# Reactive astrocytes transduce blood-brain barrier dysfunction through a TNFα-STAT3 signaling axis and secretion of alpha 1-antichymotrypsin

**DOI:** 10.1101/2022.02.21.481336

**Authors:** Hyosung Kim, Kun Leng, Jinhee Park, Alexander G. Sorets, Suil Kim, Alena Shostak, Sarah M. Sturgeon, Emma H. Neal, Douglas G. McMahon, Matthew S. Schrag, Martin Kampmann, Ethan S. Lippmann

## Abstract

Astrocytes are critical components of the neurovascular unit that support blood-brain barrier (BBB) function in brain microvascular endothelial cells (BMECs). Transformation of astrocytes to a reactive state in response to injury and disease can be protective or harmful to BBB function, but the underlying mechanisms for these effects remain mostly unclear. Using a human induced pluripotent stem cell (iPSC)-derived coculture model of BMEC-like cells and astrocytes, we found that tumor necrosis factor alpha (TNFα) transitions astrocytes to an inflammatory reactive state through activated STAT3 signaling, whereby the resultant astrocytes disrupt passive BBB function and induce vascular cell adhesion molecule 1 (VCAM-1) expression in the BMEC-like cells. These associations between inflammatory reactive astrocytes, STAT3 activation, and vascular VCAM-1 expression were corroborated in human postmortem tissue. Bioinformatic analyses coupled with CRISPR interference techniques in the iPSC model revealed that inflammatory reactive astrocytes transduce BBB disruption in part through *SERPINA3*, which encodes alpha 1-antichymotrypsin (α1ACT), a secreted serine protease inhibitor associated with aging, neuroinflammation, and Alzheimer’s disease. In murine *ex vivo* cortical explant cultures, shRNA-mediated silencing of *Serpina3n* in astrocytes reduced vascular VCAM-1 expression after TNFα challenge. Further, direct treatment with recombinant Serpina3n in both *ex vivo* explant cultures and the brain *in vivo* (via intracerebroventricular injection into wild-type mice) was sufficient to induce vascular VCAM-1 expression and reduce tight junction integrity. Overall, our results define the TNFα-STAT3 signaling axis as a driver of an inflammatory reactive astrocyte subtype responsible for BBB dysfunction. Our results also identify α1ACT as an explicit mediator of BBB damage and suggest that inhibition of α1ACT expression or activity could represent a therapeutic avenue for reversing BBB deficits in aging and neurodegenerative disease.

## Introduction

Specialized brain microvascular endothelial cells (BMECs) comprise the functional component of the blood-brain barrier (BBB), and together with supporting astrocytes, pericytes, and neurons, they collectively form the neurovascular unit (NVU). This NVU maintains the selective balance of molecules between the bloodstream and brain and its dynamic activity is essential for neural health and homeostasis^1–3^. The role of pericytes in BBB development, maintenance of healthy BBB function, and loss of BBB integrity during disease has become increasingly clear in recent years^4^. Likewise, newer studies are providing insight into how neuronal activity regulates local BBB functions^5^. In contrast, astrocytes, which were historically believed to be essential to BBB development and maintenance^6^, now have a less clear role in these activities. Seminal transplantation experiments in non-neural tissues showed that astrocytes could induce barrier properties in non-BBB endothelial cells^7, 8^, and subsequent studies have shown that astrocyte secreted factors such as sonic hedgehog and angiotensin can support BBB function^9–11^. Similarly, primary endothelial cells purified by immunopanning methods induced astrocyte precursor cells to differentiate into astrocytes via leukemia inhibitory factor (LIF) or LIF-like cytokines^12^, suggesting a close temporal and spatial correlation between endothelial and astrocyte differentiation. Yet, astrocytes arise postnatally after the initial onset of BBB properties, and there are conflicting reports as to whether astrocytes are acutely necessary to support BBB functions. Some reports show that delay or removal of astrocyte contacts with BMECs does not induce BBB disruption^13^, while a more recent study ablating astrocytes using tamoxifen-inducible cell death has indicated that astrocytes contribute to a non-redundant and crucial function in sustaining the adult BBB^14^. Hence, the role of astrocytes in BBB maintenance remains unclear.

This uncertainty extends into the role astrocytes may play in BBB disruption during disease. Recently, a subgroup of reactive astrocytes that can induce neuronal death in neurodegenerative conditions has been identified as a possible early driver of neurodegeneration. Inflammatory cytokines secreted by classically activated microglia induce the transition of normal astrocytes to this ‘inflammatory reactive’ phenotype, which is highly enriched in both aged and neuropathologic human tissues^15–17^. While inflammation caused by microglia is implicated in neurodegeneration, reactive astrocyte-associated inflammation may also play a key role. Indeed, in mouse models of amyotrophic lateral sclerosis^18^, Parkinson’s disease^19^, and white matter injury^20^, the prevention of inflammatory astrocyte conversion is neuroprotective. However, the effects of such reactive astrocytes on the BBB are unclear. Some studies have shown that reactive astrocytes in pathological conditions adversely affect endothelial integrity via secreted proteins, such as vascular endothelial growth factor (VEGF)^21–23^ and interleukin 6 (IL-6)^22^. Astrocytes have also been shown to secrete factors that mediate leukocyte recruitment^24, 25^. Likewise, diseases that exhibit increased numbers of reactive astrocytes are also typically associated with BBB disruption^26, 27^. Yet, some studies have also shown that reactive astrocytes protect BBB functions after neural injury^28–30^. Hence, a detailed understanding of the molecular connections between reactive astrocyte subtypes and BBB disruption or repair remains largely unknown. Such limitations may be due in part to the heterogeneity of reactive astrocytes *in vivo* and the challenges associated with testing how the conversion of astrocytes to a reactive state influences BBB properties.

With the remarkable progress of human induced pluripotent stem cell (iPSC) technology, differentiation protocols now facilitate access to various cell types of NVU, which offers unprecedented opportunities to study human cell biology and cell-cell interactions in health and disease^31^. Astrocytes can be efficiently derived from iPSCs by temporal application of specific signaling factors, and newer protocols can be carried out under serum-free conditions to prevent acquisition of a reactive status^32, 33^. In contrast, most protocols for generating BMEC-like cells have utilized serum-containing medium^34^, which previously prevented the study of reactive transition states in BMEC-astrocyte cocultures. To address this issue, we recently developed serum-free strategies for differentiating iPSCs to BMEC-like cells, including the use of basal media typically employed in astrocyte cultures^35, 36^. These advancements provide a highly controlled system to probe how reactive astrocyte transitions influence BBB properties in BMECs in the absence of other NVU cell types.

Here, we used iPSCs to reveal that a tumor necrosis factor alpha (TNFα)-STAT3 signaling axis generates reactive astrocytes that induce BBB dysfunction through loss of passive barrier function and upregulation of vascular cell adhesion molecule 1 (VCAM-1). We mapped astrocyte-specific pathways putatively involved in BBB disruption using RNA sequencing and various bioinformatics tools, then utilized CRISPR interference (CRISPRi) techniques to systematically silence different genes for loss-of-function studies, which identified *SERPINA3* (encoding alpha 1-antichymotrypsin, or α1ACT) as a candidate astrocyte factor causing BBB disruption. Follow-up experiments in mouse cortical explant cultures and in wild-type mice *in vivo* confirmed these associations between reactive astrocytes, α1ACT, and BBB dysfunction. Collectively, these results identify an inflammatory reactive astrocyte state responsible for negative BBB outcomes and define α1ACT as a novel mediator of this process.

## Results

### GBP2^+^ reactive astrocytes are adjacent to inflamed VCAM-1^+^ blood vessels in various human neuropathological conditions

In advance of our *in vitro* studies, we sought to confirm the association between reactive astrocytes and dysfunctional BMECs in human brain tissue. Changes to the BBB and NVU, including inflamed endothelial cells and reactive astrocytes, are a common feature related to aging and neurodegenerative diseases^37, 38^, but to our knowledge, the relative abundance and correlation of these changes have not been previously evaluated. In singular cases of patients exhibiting unspecified convulsive seizure disease and cerebral amyloid angiopathy with hemorrhage, we observed co-localization of GFAP and GBP2, a marker for inflammatory reactive astrocytes^17^, adjacent to blood vessels in the cerebral cortex (Fig. 1A-B). To more quantitatively investigate whether inflammatory reactive astrocytes are specifically adjacent to inflamed vessels, we probed tissue sections from an Alzheimer’s disease (AD) cohort and asymptomatic age-matched controls for GFAP, GBP2, and VCAM-1 expression. We inferred from morphology that the majority of VCAM-1 expression in the grey matter was localized to blood vessels in both aged individuals and AD patients (Fig. 1C). In addition, the expression of GBP2 was pronounced in subpopulations of GFAP^+^ astrocytes in both cohorts (Fig. 1C-D). Overall, the expression levels of vascular VCAM-1 and astrocytic GBP2 were higher in the AD cohort compared to the age-matched asymptomatic cohort (Fig. 1E and G-H). Since inflammatory reactive astrocytes are known to secrete harmful cytokines that contribute to neuroinflammation, we analyzed the relationship between the expression of GBP2^+^ astrocytes and VCAM-1^+^ endothelial cells in both cohorts using a Pearson correlation test (Fig. 1I-J). As expected, a positive correlation was observed in the asymptomatic cohort (p = 0.0008, Pearson r^2^ = 0.34168) and AD cohort (p = 0.0001, Pearson r^2^ = 0.52526), suggesting interactions between dysfunctional BMECs and reactive astrocytes as a function of aging and disease.

**Figure 1.**
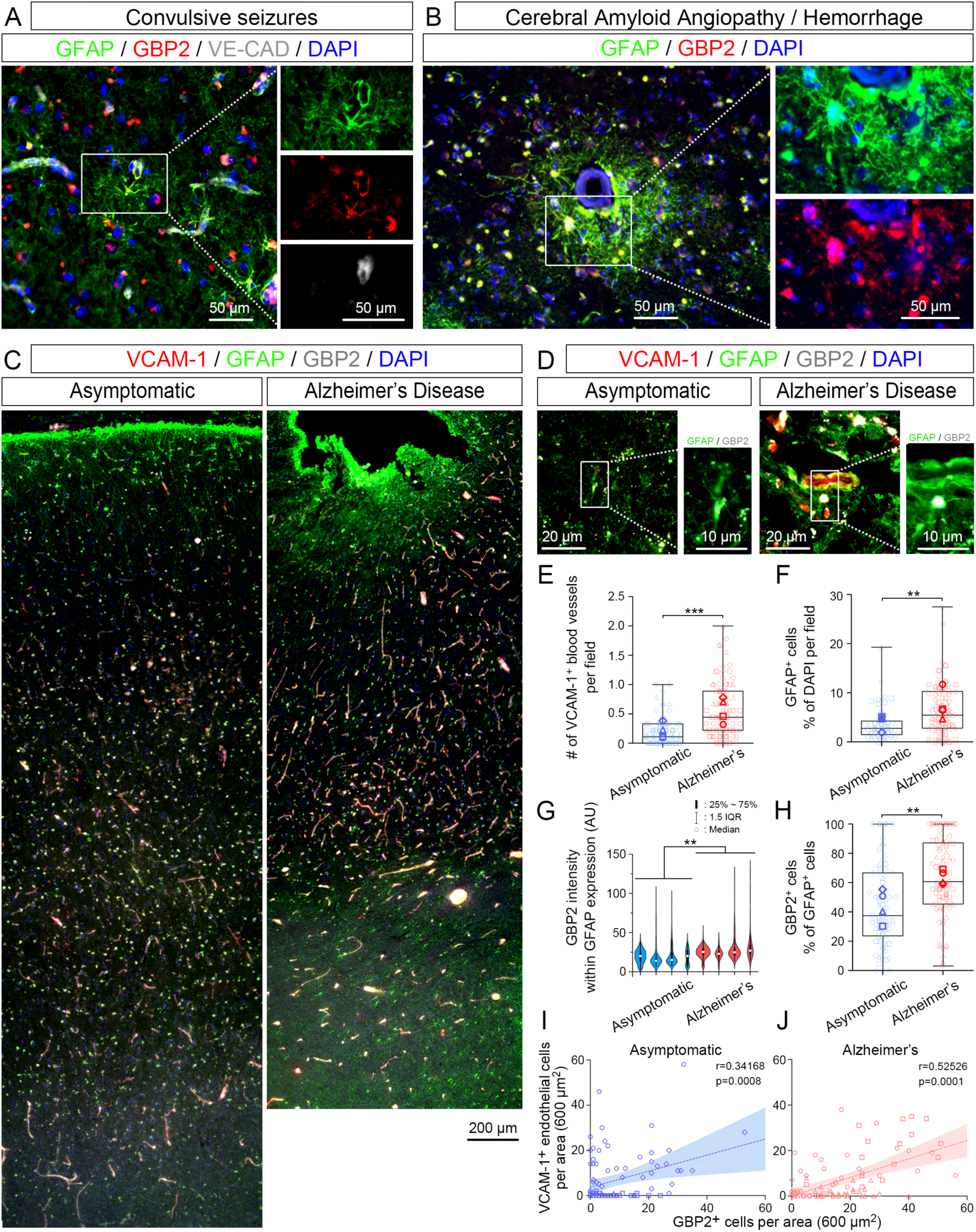
Hallmarks of GBP2^+^ inflammatory reactive astrocytes and vascular VCAM-1 expression in human brain tissue isolated from patients with neuropathological diseases. (A) Representative images of perivascular GBP2^+^/GFAP^+^ astrocytes in brain tissue from a patient with unspecified convulsive seizures. (B) Representative images of perivascular GBP2^+^/GFAP^+^ astrocytes in brain tissue from a patient with cerebral amyloid angiopathy and associated hemorrhage. (C) Representative images of VCAM-1, GFAP, and GBP2 expression across the cortex of an asymptomatic patient (left) versus a patient diagnosed with Alzheimer’s disease (right). (D) Representative cortical perivascular images of VCAM-1, GFAP, and GBP2 expression in an asymptomatic patient (left) versus a patient diagnosed with Alzheimer’s disease (right). (E-H) Quantification of VCAM-1^+^ vessels (panel E), GFAP^+^ astrocytes (panel F), GBP2 intensity in GFAP+ astrocytes (panel G), and the percentage of GBP2^+^ astrocytes (panel H) in age-matched asymptomatic brains and Alzheimer’s disease brains (n=4 biological replicates for each condition). The boxes for panels E, F, and H show the range between the 25^th^ and 75^th^ percentiles, the line within each box indicates the median, and the outer lines extend to 1.5 times the interquartile range (IQR) from the box. Faint datapoints indicate individual images and solid datapoints indicate the mean for each biological replicate. The violin plot (panel G) shows the same features for each biological replicate. Student’s t-test: **, p<0.01; ***, p<0.001. (I-J) Correlation of VCAM-1^+^ cells to GBP2^+^ cells per area in age-matched asymptomatic brains (panel I) and Alzheimer’s disease brains (panel J) (n=4 biological replicates for each condition). Best-fit lines with 95% confidence intervals from linear regression, as well as Pearson correlation coefficients and p-values, are presented on each plot.

### Soluble crosstalk between BMECs and inflammatory reactive astrocytes disrupts passive BBB function in a coculture-specific manner

To investigate direct interactions between human BMECs and reactive astrocytes, we employed serum-free iPSC systems established in previous studies. iPSCs were differentiated to CD44^+^/GFAP^+^ astrocytes by standard neuralization, manual selection of primitive neural progenitor cells, and treatment with BMP4 and FGF2 for three weeks (Fig. 2A-B). In separate cultures, iPSCs were differentiated to BMEC-like cells under serum-free conditions, where immunostaining was used to detect continuous occludin^+^ tight junctions in line with high transendothelial electrical resistance (TEER) that could be maintained above 1,000 Ω×cm^2^ for up to 2 weeks, with and without astrocyte coculture (Fig. 2C-2E). Because secreted factors derived from quiescent astrocytes in non-contact coculture systems support BBB properties^35^, we first investigated if secreted factors derived from inflammatory reactive astrocytes would negatively influence TEER within the BMEC-like cells. We confirmed that treatment of the astrocytes for 48 hours with the canonical inflammatory reactivity inducers interleukin 1 alpha (IL-1α), TNFα, and complement subcomponent q (C1q)^17^ yielded hypertrophic amoeboid shapes with GBP2 expression, indicative of transition to an inflammatory reactive state (Fig. 2F). Next, astrocytes were maintained in serum-free medium for 72 hours with or without IL-1α, TNFα, and C1q, and the spent medium was collected, concentrated, and fed to monocultured BMEC-like cells (Fig. 2G). Surprisingly, no significant differences in TEER were observed (Fig. 2H), suggesting active crosstalk might be necessary to transduce the damaging effects of inflammatory reactive astrocytes. To probe this possibility, we generated monocultures of BMEC-like cells and cocultures of BMEC-like cells with astrocytes, then added single doses of various inflammatory cytokines (Fig. 2I). As expected from previous work (Fig. 2H)^39, 40^, BMEC-like cells in monoculture were unresponsive to cytokine treatments. In contrast, in the coculture system, treatment with TNFα or IL-1α was sufficient to significantly reduce TEER in the BMEC-like cells, whereas C1q and interferon gamma (IFNγ) treatments did not yield BBB disruption, highlighting cytokine-specific responses (Fig. 2J). Interestingly, we observed a lag time of several days between the addition of cytokines and decreased TEER, suggesting other factors beyond the cytokines were contributing to BBB dysfunction. Hence, we next sought to analyze the changes to each cell population at later time points.

**Figure 2.**
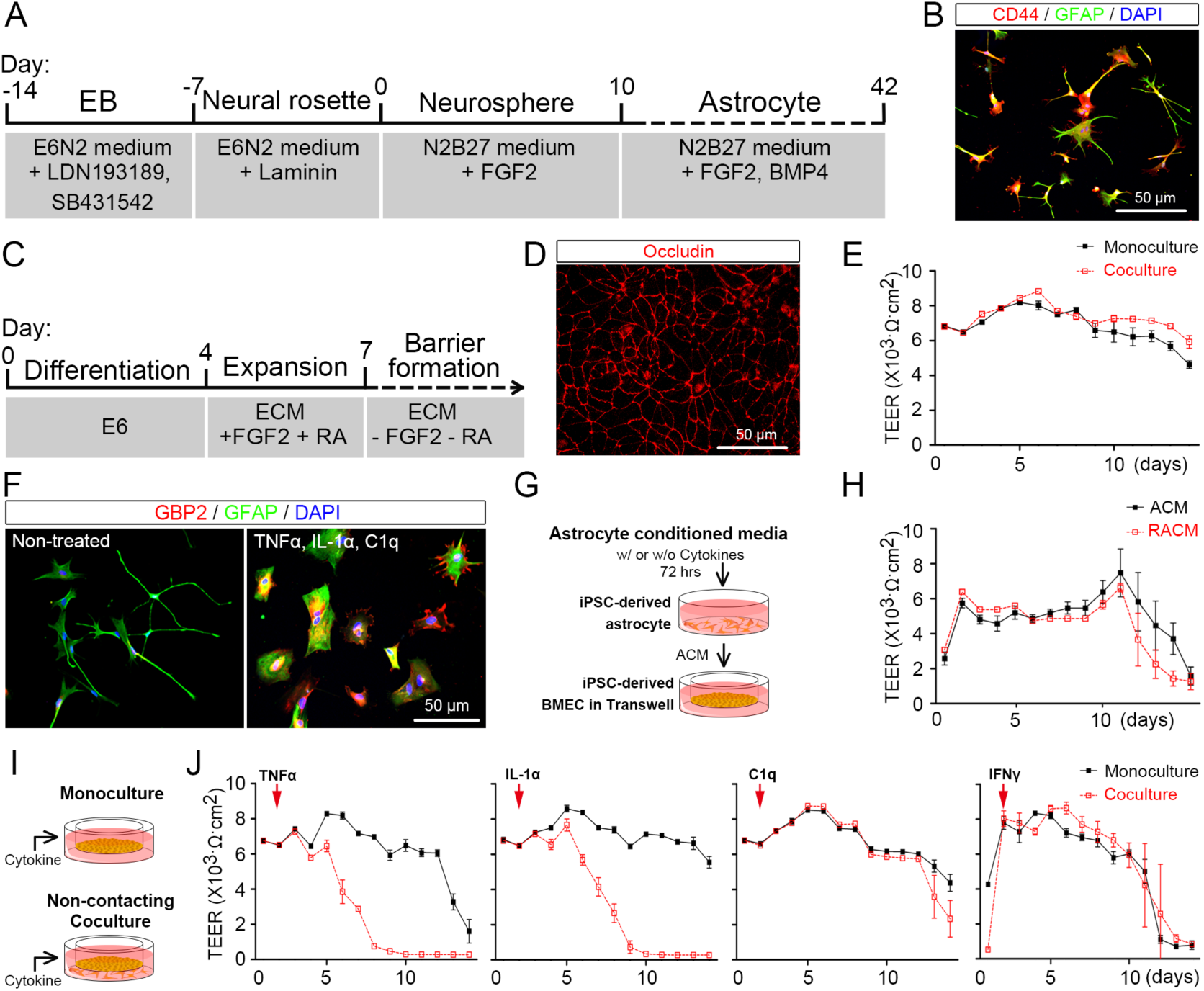
Coculture of iPSC-derived BMEC-like cells and astrocytes is necessary for disruption of passive barrier function by inflammatory cytokines. (A) Schematic procedure for deriving astrocytes from iPSCs. (B) Representative immunofluorescent images of astrocyte markers after 40 days of differentiation. (C) Schematic procedure for deriving BMEC-like cells from iPSCs. (D) Representative immunofluorescent images of smooth and continuous tight junctions in the BMEC-like cells 8 days after differentiation. (E) Representative TEER values in BMEC-like cells in monoculture or in coculture with astrocytes. Trends were confirmed across biological n=3. (F) Representative immunofluorescent images of GFAP and GBP2 expression in astrocytes 48 hours after treatment with an inflammatory cytokine cocktail. (G-H) Schematic diagram of the procedure for treating BMEC-like cells with astrocyte conditioned medium (panel G) and subsequent TEER measurements (panel H). Astrocyte conditioned medium without inflammatory cytokines is noted as “ACM” and astrocyte conditioned medium collected after cytokine treatment is referred to as reactive astrocyte conditioned medium, or “RACM.” Trends were confirmed across biological n=3. (I-J) Experimental setup (panel I) and subsequent TEER measurements (panel J) after dosing BMEC-like cells with cytokines in monoculture or coculture. Trends were confirmed across biological n=3.

### Transcriptome profiling reveals early-stage inflammatory interactions between BMECs and astrocytes

To gain mechanistic insight into how reactive astrocytes contribute to BBB dysfunction in the *in vitro* model, we constructed, mapped, and compared differences of transcriptomes and gene regulatory networks using bulk RNA sequencing (RNA-seq). First, to find an appropriate concentration of TNFα for studying astrocyte/BMEC interactions, we measured TEER in response to doses of TNFα ranging from 1 ng/ml to 100 ng/ml. Single TNFα doses of 10 and 100 ng/ml were sufficient to cause a decrease in TEER and an increase in *VCAM1* expression in BMEC-like cells (Supplementary Fig. 1). Based on these experiments, we proceeded with a 10 ng/ml dose of TNFα for transcriptional analyses, specifically isolating RNA at day 5 of culture at the time point when TEER started to decline in the coculture condition. This time point was also used for cells in monoculture (BMEC-like cells or astrocytes) with or without TNFα treatment. First, in bulk RNA-seq datasets, we checked the cellular identity of astrocytes and endothelial cells in the absence of TNFα to further validate the *in vitro* system composed of iPSC-derived cells. In accordance with key functional properties of BBB such as elevated TEER and continuous occludin^+^ tight junctions shown in Figure 2D-E, we could detect endothelium- and BBB-specific gene expression including *CDH5* (VE-cadherin), *TEK* (Tie2), *VWF* (von Willebrand factor), and *SLC2A1* (GLUT1) in the iPSC-derived BMEC-like cells (Supplementary Fig. 2A). Likewise, astrocytic identity was confirmed by astrocyte-specific markers including *GFAP*, *CD44*, *SLC1A2* (GLT-1), and *AQP4* (Aquaporin-4) (Supplementary Fig. 2B). To obtain a general picture of the relationship among different culture conditions, we performed unbiased dimensionality reduction with all detected transcripts using principal-component analysis (PCA) and visualized the relationship with the first two principal components. PCA on BMEC-like cell transcriptomes showed two distinct clusters defined by TNFα, but not by culture condition (monoculture or coculture with astrocytes; Fig. 3A). Volcano plots and gene ontology (GO) analyses on differentially expressed genes (DEGs) were also constructed to analyze TNFα effects on monocultured versus cocultured BMEC-like cells (Fig. 3B-E). Among the DEGs in BMEC-like cells, adhesion molecules including *VCAM1*, *CEACAM1*, and *ICAM1* were substantially induced by TNFα in both monoculture and coculture, whereas junctional adhesion molecule 1 (*F11R*) was downregulated in both monoculture and coculture. Major genes involved in the formation and maintenance of adherence junctions, *CDH5* and *CADM3,* were also downregulated by TNFα regardless of the culture system. In contrast, TNFα induced the upregulation of *CD47*, an integrin-associated protein involved in age-associated deterioration in angiogenesis^41^, only in the coculture system. Key endocytosis molecules were downregulated by TNFα in both culture conditions, for instance, *PACSIN3* (important regulator of internalization of plasma membrane proteins^42^) and *RAB5C* (a small GTPases key regulator of trafficking of integrin complex in endothelial cells^43^). Exposure to TNFα also caused the downregulation of the proangiogenic genes such as *AGGF1*, *FGF2*, and *ANG*^44–46^. Further, while many overlapping DEGs could be identified, we observed key differences in gene ontology. Notably, in the coculture condition, inflammatory signaling pathways, including “response to tumor necrosis factor,” “STAT cascade,” and “neuroinflammatory response” were selectively enriched (Fig. 3D-E).

**Figure 3.**
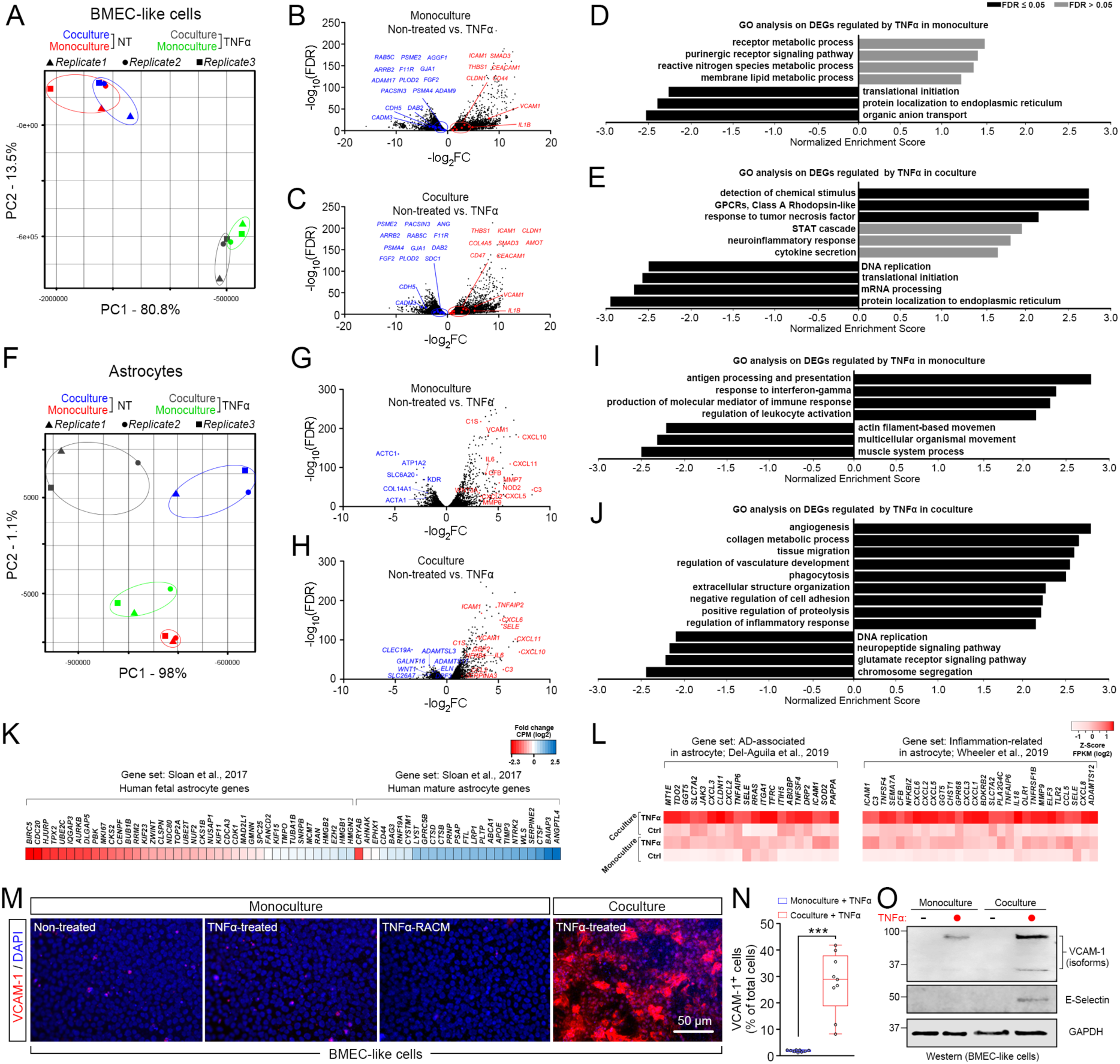
Comparison of transcriptomic signatures and inflammatory phenotypes in astrocytes and BMEC-like cells as a function of coculture and treatment with TNFα. (A) PCA clustering plot of RNA-seq data in BMEC-like cells from four different conditions with n=3 biological replicates per condition. (B-C) Volcano plot of DEGs (FDR<0.1) in BMEC-like cells in monoculture versus coculture, with or without TNFα treatment. Red and blue points highlight genes associated with endothelial cell-cell junctions, internalization and trafficking, and angiogenesis. (D-E) GO analyses in the BMEC-like cells in monoculture versus coculture after TNFα treatment. Normalized enrichment scores and false discovery rate are provided. (F) PCA clustering plot of RNA-seq data in astrocytes from four different conditions and n=3 biological replicates per condition. (G-H) Volcano plot of DEGs (FDR<0.1) in astrocytes in monoculture versus coculture, with or without TNFα treatment. Red and blue points highlight genes associated with inflammation, chemokines, and immune responses. (I-J) Select GO analyses in the astrocytes in monoculture versus coculture after TNFα treatment. Normalized enrichment scores and false discovery rate are provided. (K) Heat map showing fold change of gene expression of maturity-related genes in cocultured astrocytes compared to monocultured astrocytes. The gene set is based on Sloan et al., 2017^47^. The color intensity of the heat map represents the log2(CPM value) for each gene. (L) Heat map showing relative expression of genes associated with AD and inflammation across all astrocyte conditions. The gene sets are based on Del-Aguila et al., 2019^48^ (left panel) and Wheeler et al., 2019^103^ (right panel). The color of the heat map represents the log2(FPKM value) for each gene. (M-N) Representative immunofluorescent images of VCAM-1 expression in BMEC-like cells under different conditions (panel M) and select quantification (panel N). Data were acquired at day 14 according to the timing presented in Figure 2J. Quantification was conducted using n=3 biological replicates. The box shows the range between the 25th and 75th percentiles, the line within each box indicates the median, the outer lines extend to 1.5 times the interquartile range from the box, and each data point represents an individual image (3 images quantified per biological replicate). Student’s t-test: ***, p<0.001. (O) Representative western blot showing relative VCAM-1 and E-selectin expression in BMEC-like cells in monoculture or coculture, with or without TNFα treatment. Data were acquired at day 14 according to the timing presented in Figure 2J. GAPDH was used as a loading control. Trends were confirmed across n=3 biological replicates.

Meanwhile, the PCA on astrocytes showed both coculture- and TNFα-responsive transcriptomic signatures. The first principal component (PC1) was dominated by TNFα, and the second principal component (PC2) separated the samples according to culture condition (Fig. 3F). Astrocytes treated with TNFα in monoculture upregulated many inflammation-associated genes with concurrent enrichment of inflammatory signaling pathways (Fig. 3G-I), and many of these astrocytic genes were also upregulated in the TNFα-treated coculture condition (Fig. 3H-J), but some different enriched pathways associated with vascular interactions were observed, including “angiogenesis” and “regulation of vasculature development.” Based on these results, we compared the gene expression signatures in astrocytes to several published datasets. In astrocytes not treated with TNFα, we observed a profound shift in gene expression towards a mature signature if astrocytes were cocultured with BMEC-like cells (Fig. 3K)^47^.

Likewise, compared to monoculture astrocytes treated with TNFα, coculture with BMEC-like cells amplified expression of inflammation- and AD-associated genes (Fig. 3L)^48^. These results suggest that iPSC-derived astrocytes acquire a more mature profile upon coculture with BMEC-like cells, which in turn may facilitate their inflammatory responses to TNFα.

The enhanced astrocytic inflammation in the coculture condition led us to further analyze inflammatory phenotypes in the BMEC-like cells. A key molecular phenotype of endothelial cells in response to inflammation is the expression of adhesion molecules and selectins^49^. Here, we primarily focused on VCAM-1, which has recently been implicated in neurodegeneration^50^ as well as our human tissue analyses in Figure 1. Although previous studies with human endothelial cell lines showed the prompt expression of VCAM-1 following stimulation with inflammatory cytokines^51^, a lack of adhesion molecule expression in response to inflammation has been noted as a specific shortcoming in many iPSC-derived BBB models. As described previously, in the iPSC-derived BMEC-like cells (cocultured with astrocytes), 10 and 100 ng/ml TNFα increased *VCAM1* by 2- and 4-fold, respectively, 2 weeks after coculture in the presence of TNFα (Supplementary Fig. 1B), which mirrored a decline in TEER (Fig. 2J). We therefore examined VCAM-1 protein expression on the iPSC-derived BMEC-like cells under various conditions. Immunostaining revealed minimal VCAM-1 expression in monoculture BMEC-like cells with or without TNFα treatment, despite strong induction of *VCAM1* in response to TNFα. Conditioned medium from TNFα-treated reactive astrocytes was also unable to induce VCAM-1 expression, again indicating the importance of astrocyte-endothelial crosstalk in the inflammatory response. In contrast, in the coculture system, TNFα treatment induced widespread VCAM-1 expression (Fig. 3M-N). The effects of coculture were confirmed by western blotting, where we also determined that E-Selectin, another key adhesion molecule, was solely induced by TNFα in the coculture condition (Fig. 3O). Overall, the results from these experiments highlight the importance of astrocyte-endothelial crosstalk for transducing inflammatory phenotypes in the *in vitro* BBB model, potentially through post-transcriptional mechanisms.

### TNFα-mediated BBB disruption is dependent on astrocytic STAT3 signaling in the *in vitro* coculture system

We next focused on the inflammatory signaling axis that could be responsible of TNFα-mediated BBB disruption. Since BBB dysfunction in our *in vitro* model was transduced by inflammatory reactive astrocytes in a non-contacting coculture setup (Fig. 2J and Fig. 3M-O), we first focused on analyzing soluble factors secreted by astrocytes. To predict BBB-disruptive and immune-stimulative soluble factors provided by reactive astrocytes, we organized reactive astrocytic transcripts based on a database composed of 6,943 secreted human proteins^52^ and then computationally predicted interactions with endothelial receptome genes (Fig. 4A and Supplementary Fig. 3). Within the astrocytes, we generally observed enriched transcripts for target genes along the NF-κB/STAT3 axis that were enhanced by TNFα treatment; these astrocyte genes included *CCL2*, *CXCL10*, *CXCL11*, *CXCL2*, *CXCL3*, *CXCL8*, *EDN1,* and *SERPINA3*^53–58^. We also observed enriched mapping of astrocyte-secreted factors and endothelial receptors for complement genes, including astrocytic upregulation of *C1S*, *C1RL*, *C3*, and *CFB*. Furthermore, using the Search Tool for the Retrieval of Interacting Genes/Proteins (STRING)^59^ database decoyed by STAT3, we found that 22 out of the top 50 astrocytic secretome genes were closely correlated with STAT3 (Fig. 4B). In addition, an Ingenuity Pathway Analysis conducted on DEGs predicted that NF-κB and STAT3 signaling were preferentially activated by TNFα treatment in coculture compared to monoculture (Fig. 4C).

**Figure 4.**
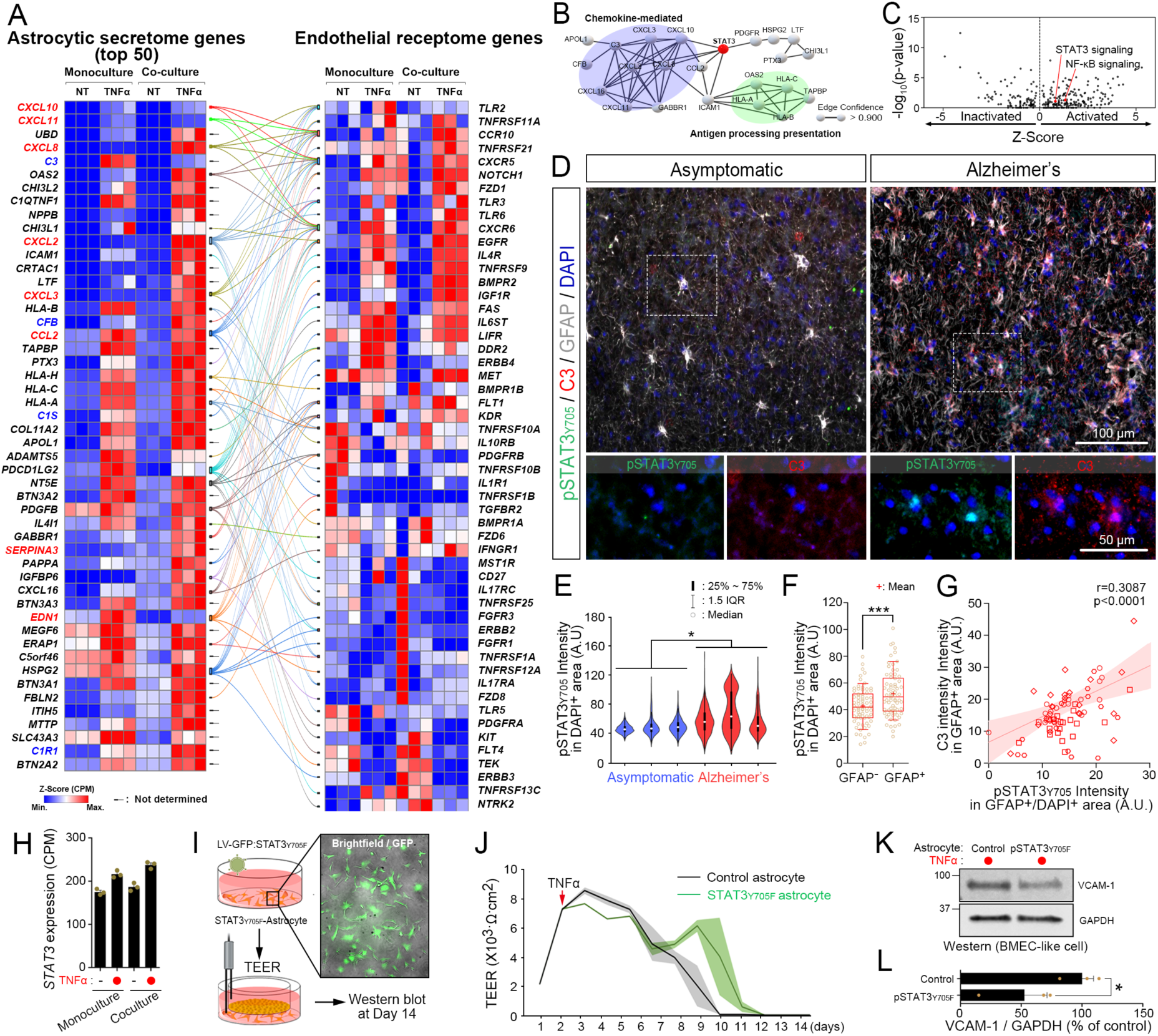
STAT3 signaling is activated in reactive inflammatory astrocytes and directly influences BBB dysfunction. (A) Network analysis (confidence score >0.7) of the top 50 differentially expressed astrocytic genes encoding secreted factors, with corresponding endothelial genes encoding receptors. The network was produced by STRING and visualized by Origin 2021. Links are color-coded to the astrocytic source node. (B) Network analysis (confidence score >0.9) of the STAT3 interactome suggests involvement of 22 out of the top 50 astrocytic genes encoding secreted factors. The STAT3 bait protein is labeled in red. Chemokine-mediated genes and antigen processing presentation-related genes are circled in purple and green, respectively. (C) Ingenuity Pathway Analysis predicts activation of STAT3 and NF-kB signaling in TNFα-treated astrocytes in coculture with BMEC-like cells. Activities were calculated by importing DEGs (FDR<0.1) between cocultured astrocytes with or without TNFα treatment. (D) Representative images of pSTAT3_Y705_, C3, and GFAP expression in the cortex of an asymptomatic patient (left) versus a patient diagnosed with Alzheimer’s disease (right). (E) Quantification of nuclear pSTAT3_Y705_ intensity in the cortex of asymptomatic patients versus patients diagnosed with Alzheimer’s disease (758-883 cells counted per sample; n=3 biological replicates). Violin plots show the range between the 25^th^ and 75^th^ percentiles, the line within each box indicates the median, and the outer lines extend to 1.5 times the IQR from the box. Student’s t-test: *, p<0.05. (F) Quantification of nuclear pSTAT3_Y705_ intensity in GFAP^-^ versus GFAP^+^ cells in the cortex of patients diagnosed with Alzheimer’s disease (70 cells counted per sample; n=4 biological replicates). The boxes show the range between the 25^th^ and 75^th^ percentiles, the line within each box indicates the median, the plus sign indicates the mean, and the outer lines extend to 1.5 times the IQR from the box. Student’s t-test: ***, p<0.001. (G) Correlation of C3 intensity in GFAP^+^ astrocytes to nuclear pSTAT3_Y705_ intensity of GFAP^+^ astrocyte in the cortex of four patients diagnosed with Alzheimer’s disease. Best-fit lines with 95% confidence intervals from linear regression, as well as Pearson correlation coefficient and p-value, are presented in the plot. (H) Bar graph showing STAT3 gene expression (CPM) in astrocytes at day 5 of monoculture or coculture, with or without TNFα treatment. Data are graphed as mean ± SEM from n=3 biological replicates. (I) Schematic diagram of the procedure for producing dominant-negative STAT3 astrocytes for coculture with BMEC-like cells and subsequent TEER measurements and western blot analysis. (J) TEER values in BMEC-like cells in coculture with astrocytes after dominant negative STAT3 overexpression. Data are presented as a continuous mean ± shaded SEMs per condition from a single biological replicate. Trends were confirmed across n=3 biological replicates. (K-L) Representative western blot and quantification of VCAM-1 expression in BMEC-like cells. Data were acquired at day 14 according to the timing presented in Figure 2J. GAPDH was used as a loading control. Data are graphed as mean ± SEM from n=3 biological replicates. Student’s t-test: *, p <0.05.

It is well established that the STAT3 pathway mediates astrocyte reactivity in animal models^60–62^, but fewer studies have been performed in pathological human brain samples. Typically, STAT3 activation is initiated by phosphorylation on a critical tyrosine residue (Tyr705; pSTAT3), whereby pSTAT3 is then translocated to the nucleus to activate target gene expression. We immunostained tissue sections from the cortex of AD patients and asymptomatic age-matched controls to determine localization of pSTAT3_Y705_ within GFAP+ astrocytes. We further labeled for C3, a marker for reactive inflammatory astrocytes, to align with the signature of the iPSC-derived reactive astrocytes after treatment with TNFα.

We observed pSTAT3_Y705_ signal within the nuclei of GFAP^+^ astrocytes in the AD tissue, while pSTAT3_Y705_ in the asymptomatic tissue was generally situated around the nucleus (Fig. 4D-F and Supplementary Fig 4). Nuclear pSTAT3_Y705_ was also positively correlated with C3 expression (Fig. 4G), highlighting the connections between the *in vitro* coculture model and human neuropathology.

Based on these data, we revisited the *in vitro* coculture model to further clarify the connections between astrocytic STAT3 activation and BBB dysfunction. Similar levels of astrocytic *STAT3* expression were observed under all culture conditions (Fig. 4H), so we decided to inactivate STAT3 signaling in astrocytes by constitutively overexpressing human Y705F dominant-negative STAT3 using lentiviral transduction^63^ (Fig. 4I), which significantly reduced VCAM-1 expression and extend TEER longevity in BMEC-like cells after TNFα treatment relative to astrocytes transduced with the empty lentiviral backbone (Fig. 4J-L). Since VEGF secretion by astrocytes has been tied to BBB disruption in the inflammatory experimental autoimmune encephalomyelitis (EAE) mouse model^21, 64^, we also queried the role of VEGF in the *in vitro* model. We found that *VEGFA* expression was mostly unchanged by TNFα treatment, and knockdown of *VEGFA* in astrocytes in the TNFα-treated coculture system using a CRISPR interference strategy (CRISPRi, described in more detail below) did not reduce VCAM-1 expression or alter TEER in BMEC-like cells (Supplementary Fig. 5). Hence, the TNFα-STAT3 signaling axis in inflammatory reactive astrocytes contributes to BBB dysfunction, but likely not through VEGF secretion.

### TNFα-stimulated reactive astrocytes yield BBB dysfunction through expression of *SERPINA*3

To further elucidate astrocytic factors that could transduce BMEC inflammation and cause BBB disruption, we used the STRING tool to cluster genes for easier interrogation. The STRING analysis identified 11 genes that were predominantly upregulated in cocultured astrocytes stimulated with TNFα, and these genes were segregated into 3 clusters roughly based on their known functions (Fig. 5A-B). We developed pooled sgRNAs targeting each cluster, which were transduced into CRISPRi astrocytes by lentivirus. In this strategy, iPSCs were modified to express dead Cas9 fused to the KRAB repressor domain and differentiated to astrocytes before receiving the lentivirus. These transduced astrocytes were then cocultured with BMEC-like cells and treated with TNFα, followed by assessments of TEER and VCAM-1 expression (Fig. 5C). Suppression of the chemokine gene cluster (*CXCL10*, *CXCL8*, *CXCL6*, and *CCL2*) did not affect TEER or VCAM-1 expression, which was potentially expected since these genes are mainly involved in recruiting and activating neutrophils^65, 66^. In contrast, suppression of the *SOD2*, *DSP*, and *JAK3* cluster increased VCAM-1 expression without affecting TEER. These data are consistent with a previous observation that decreasing SOD2 by TFEB knockdown induced VCAM-1 expression in non-brain endothelial cells^67^. Furthermore, TNFα stimulation of STAT3 signaling in astrocytes is typically associated with JAK1 or JAK2 but not JAK3^68, 69^, also consistent with our data. In contrast, suppression of *TNFAIP2*, *C3*, *SLC43A2*, and *SERPINA3*, all of which are suggested to be activated by STAT3^58, 70–72^, decreased VCAM-1 expression and increased TEER longevity in BMEC-like cells (Fig. 5D-F), which is consistent with the results observed after STAT3 signaling inactivation in astrocytes. Based on these results, we silenced the individual genes from this cluster and re-assessed VCAM-1 expression, which revealed *SERPINA3* as a key effector for mediating BBB damage (Fig. 5G-H). Since C3 has been previously identified as an astrocyte-derived factor that causes BBB inflammation in mouse models of aging and tauopathy^73^, and we observed a trending but not statistically significant reduction in VCAM-1 expression from *C3* knockdown, we subsequently assessed the effects of simultaneous *SERPINA3* and *C3* knockdown. Here, knockdown of both genes yielded a similar VCAM-1 reduction compared to the individual knockdowns and a modest extension of TEER longevity (Supplementary Fig. 6), suggesting potential overlap in how these factors cause BBB damage.

**Figure 5.**
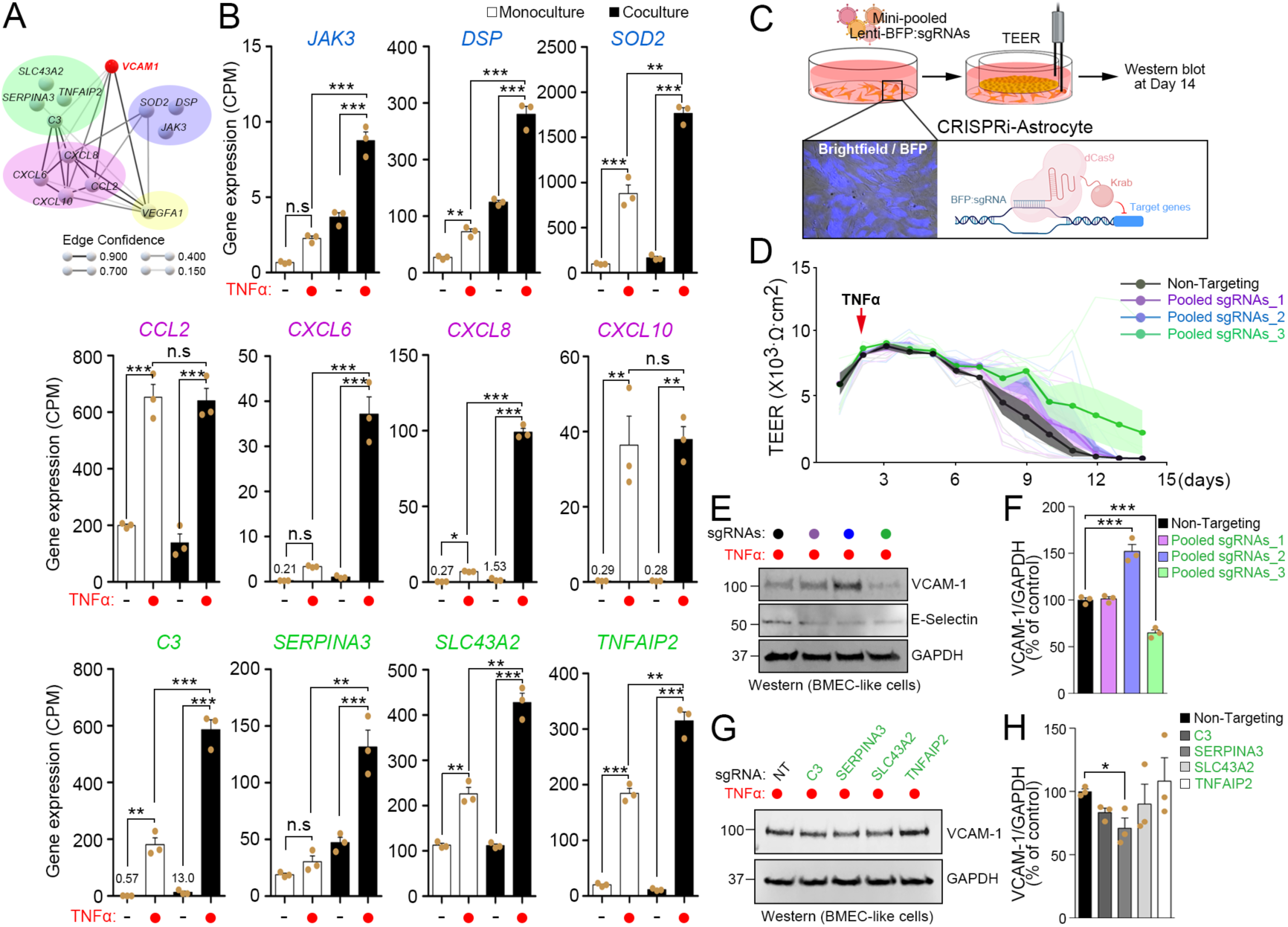
Pooled CRISPRi screen identifies *SERPINA3* expression in astrocytes as a contributor to BBB dysfunction. (A) Network representation connecting *VCAM1* to astrocytic genes predominantly upregulated by TNFα in coculture. Line thickness indicates the edge confidence. The *VCAM1* bait gene is labeled in red. Genes were segregated with colored circles into 4 clusters roughly based on their known functions. (B) Expression levels (CPM) of the selected astrocytic genes of panel A. Data are graphed as mean ± SEM from n=3 biological replicates. One-way ANOVA with Tukey’s post hoc test: *, p<0.05; **, p<0.01; ***, p<0.001. (C) Schematic procedure for pooled gene knockdown in CRISPRi-astrocytes and coculture with BMEC-like cells, followed by TEER measurements and western blot analysis. (D) TEER values in BMEC-like cells in coculture with astrocytes transduced with pooled sgRNAs. Data are presented as continuous means ± shaded SEMs aggregated from n=3 biological replicates per condition. (E-F) Representative western blot and quantification of VCAM-1 expression in BMEC-like cells cocultured with CRISPRi-astrocytes. CRISPRi-astrocytes were transduced with pooled sgRNAs targeting genes color coded to the clusters in panel A. Data were acquired at day 14 according to the timing presented in Figure 5D. GAPDH was used as a loading control. Data are graphed as mean ± SEM from n=3 biological replicates. One-way ANOVA with Tukey’s post hoc test: ***, p<0.001. (G-H) Representative western blot and quantification of VCAM-1 expression in BMEC-like cells cocultured with CRISPRi-astrocytes. CRISPRi-astrocytes were transduced with a single sgRNA targeting individual genes in the cluster colored in green in panel A. Data were acquired at day 14 according to the timing presented in Figure 5D. GAPDH was used as a loading control. Data are graphed as mean ± SEM from n=3 biological replicates. One-way ANOVA with Tukey’s post hoc test: *, p<0.05.

### Serpina3n is directly responsible for BBB dysfunction in cortical explant cultures

Serpina3n (mouse ortholog of α1ACT) has been previously identified as a reactive astrocyte marker after lipopolysaccharide treatment or middle cerebral artery occlusion^74^, but to our knowledge, no previous studies have linked α1ACT or Serpina3n to BBB dysfunction. To validate the contribution of α1ACT to BBB dysfunction, we transitioned to a mouse cortical explant model (Fig. 6A). First, to permit control over *Serpina3n* expression in astrocytes, we generated an adenovirus associated virus (AAV)-based system, in which a short GFAP (GfaBC_1_D) promoter drives expression of both EGFP and a miR-30-based short hairpin RNA^75^ (shRNAmiR; Fig. 6B). Astrocyte-specific transgene expression was validated in iPSC-derived astrocytes and mouse cortical explant cultures transduced by the AAV PHP.B capsid^76^ harboring EGFP and shRNAmiR, which demonstrated EGFP expression only to the post-stained GFAP^+^ cells; in contrast, HEK293T cells transduced by the AAV showed no EGFP expression (Fig. 6C). Next, we validated the knockdown efficiency in the explant cultures with AAV encoding an shRNA against *Serpina3n* (sh*Serpina3n*miR) or a non-targeting control shRNA (sh*Control*miR). Following TNFα treatment, EGFP^+^/GFAP^+^ astrocytes in the sh*Control*miR condition exhibited extensive Serpina3n^+^ puncta, and these puncta were significantly reduced in the sh*Serpina3n*miR condition (Fig. 6D-F). Since Serpina3n is secreted from cells, we additionally performed an ELISA on media collected from each explant culture to validate that silencing *Serpina3n* in astrocytes resulted in depletion of the soluble protein (Fig. 6G). Bolstered by this result, we transduced explant cultures with each AAV and measured VCAM-1 intensity in CD31^+^ vessels, which revealed a significant decrease in VCAM-1 in cultures transduced with sh*Serpina3n*miR relative to sh*Control*miR (Fig. 6H-I). We also treated explant cultures with recombinant mouse Serpina3n, which yielded a dose-dependent increase in VCAM-1 expression within CD31^+^ vasculature, as well as a dose-dependent decrease in claudin-5 tight junction expression within CD31^+^ vasculature and an overall reduction in vascular density (Fig. 6J-M). These results validate outcomes from the *in vitro* model and provide strong evidence that astrocyte-derived Serpina3n directly causes neurovascular inflammation and BBB dysfunction.

**Figure 6.**
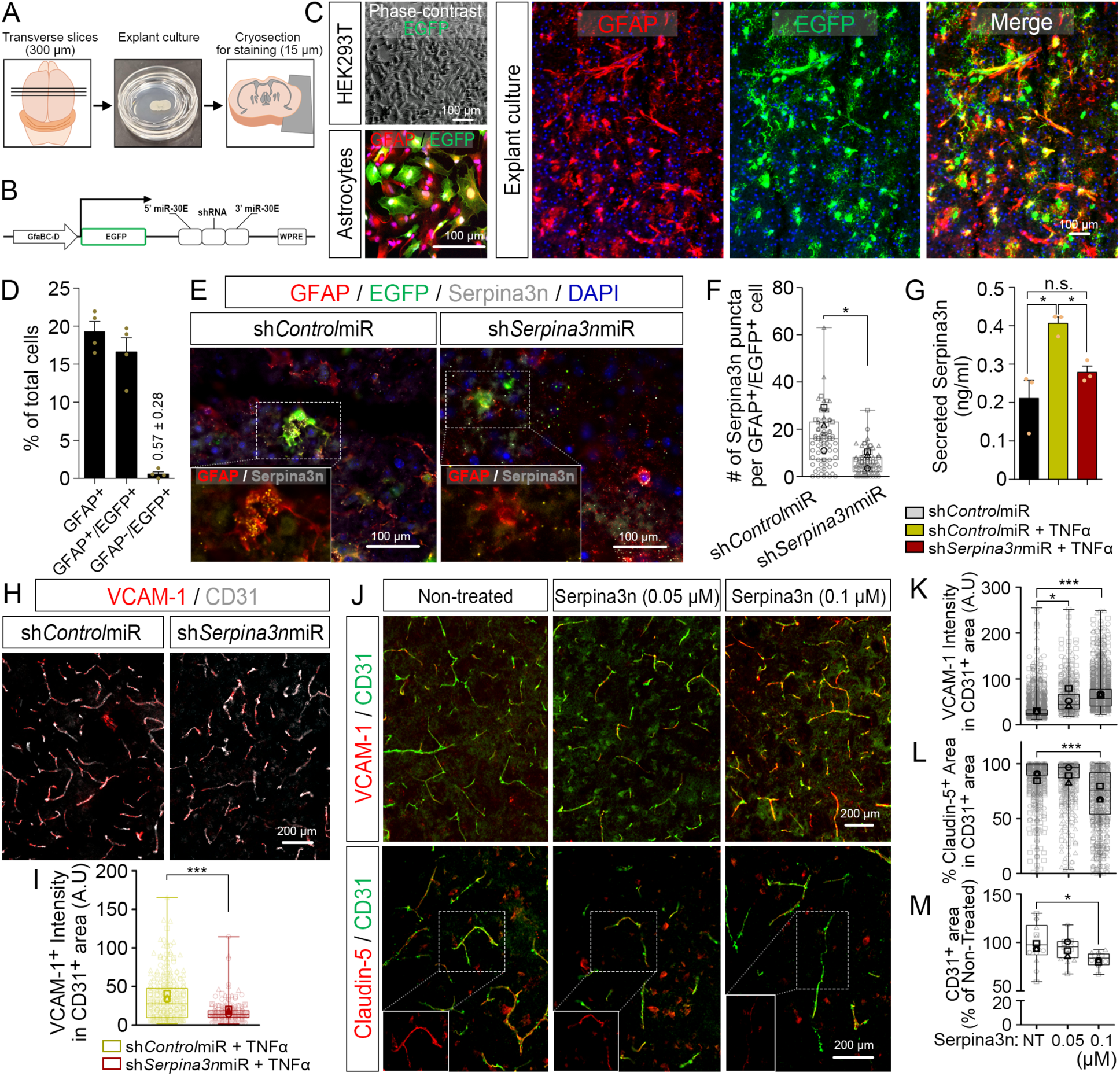
Loss- and gain-of-function experiments in cortical explant cultures connect soluble Serpina3n directly to BBB dysfunction. For all panels involving AAV transduction, cortical explants were treated with AAV for 3 days, then treated with TNFα for 3 days before fixation and analysis. For all panels involving soluble Serpina3n treatment, the protein was added for 4 days (in the absence of TNFα), followed by fixation and analysis. (A) Schematic overview of the cortical explant culture preparation and analyses. (B) Schematic representation of the master AAV vector encoding EGFP and miR-30-shRNA driven by a shortened GFAP promoter. (C) Representative images of EGFP and GFAP expression in HEK293T cells, iPSC-derived astrocytes, and cortical explant cultures after transduction with the AAV vector. (D) Quantification of EGFP and GFAP labeling within the explant cultures. Data are plotted as mean ± SEM for n=4 biological replicates. (E) Representative images of GFAP, EGFP, and Serpina3n expression in cortical explant cultures transduced with the AAV vector encoding either sh*Control*miR or sh*Serpina3n*miR. (F) Quantification of Serpina3n^+^ puncta per GFAP^+^/EGFP^+^ cell from the images in panel E. For each condition, at least 80 EGFP+/GFAP+ cells were analyzed. Each quantified cell is represented by a faint data point, and solid datapoints indicate the mean for each biological replicate (n=3). The boxes show the range between the 25^th^ and 75^th^ percentiles, the line within each box indicates the median, and the outer lines extend to 1.5 times the IQR from the box. Student’s t-test: *, p<0.05. (G) ELISA quantification of soluble Serpina3n levels in cortical explant cultures after AAV and TNFα treatment. Data are graphed as mean ± SEM from n=3 biological replicates. One-way ANOVA with Turkey’s post hoc test: n.s., p>0.05; *, p<0.05. (H) Representative images of CD31 and VCAM-1 expression in cortical explant cultures transduced with the AAV vector encoding either sh*Control*miR or sh*Serpina3n*miR. (I) Quantification of VCAM-1 intensity across total CD31^+^ area (231 vessel segments analyzed in the sh*Control*miR condition and 330 vessel segments analyzed in the sh*Serpina3n*miR condition; n=3 biological replicates per condition). The boxes show the range between the 25^th^ and 75^th^ percentiles, the line within each box indicates the median, and the outer lines extend to 1.5 times the IQR from the box. Faint datapoints indicate individual vessel segments and solid datapoints indicate the mean for each biological replicate. Student’s t-test: ***, p<0.001. (J) Representative images of endothelial VCAM-1 expression (upper panel) and claudin-5 expression (lower panel) in cortical explant cultures treated with varying concentrations of recombinant Serpina3n. (K) Quantification of VCAM-1 intensity across total CD31^+^ area. The boxes show the range between the 25^th^ and 75^th^ percentiles, the line within each box indicates the median, and the outer lines extend to 1.5 times the IQR from the box. Faint datapoints indicate individual vessel segments and solid datapoints indicate the mean for each biological replicate. One-way ANOVA with Turkey’s post hoc test: ***, p<0.001. (L) Quantification of percent claudin-5 coverage within CD31^+^ vessels (0 μM Serpina3n, 1181 vessel segments counted; 0.05 μM Serpina3n, 735 vessel segments counted; 0.1 μM Serpina3n, 673 vessel segments counted; n=3 biological replicates). The boxes show the range between the 25^th^ and 75^th^ percentiles, the line within each box indicates the median, and the outer lines extend to 1.5 times the IQR from the box. Faint datapoints indicate individual vessel segments and solid datapoints indicate the mean for each biological replicate. One-way ANOVA with Turkey’s post hoc test: ***, p<0.001. (M) Quantification of vessel density represented as the area of CD31^+^ vessels against total tissue area. Data are normalized to the non-treated control. The boxes show the range between the 25^th^ and 75^th^ percentiles, the line within each box indicates the median, and the outer lines extend to 1.5 times the IQR from the box. Faint datapoints indicate individual images and solid datapoints indicate the mean for each biological replicate. One-way ANOVA with Turkey’s post hoc test: *, p<0.05.

### Serpina3n causes BBB dysfunction after direct administration to the brain *in vivo*

Last, we investigated whether Serpina3n could directly induce BBB dysfunction *in vivo*. To better isolate its effects within the brain, recombinant mouse Serpina3n was administered by intracerebroventricular (ICV) injection. Here, a single dose of Serpina3n was sufficient to induce a significant increase in VCAM-1 expression within CD31^+^ vasculature, relative to PBS vehicle control (Fig. 7A-B). This upregulation in VCAM-1 was widespread, suggesting the effects of Serpina3n are not restricted to a specific brain region, which is consistent with a recent report showing that Serpina3n is ubiquitously induced in reactive astrocytes irrespective of location^77^. In agreement with slice culture experiments, the expression of claudin-5 was also decreased within CD31^+^ vasculature after Serpina3n injection (Fig. 7D-E). We also noted a reduction of vasculature density due to Serpina3n treatment, as measured by total area of CD31^+^ and Glut-1^+^ blood vessels (Fig. 7C and 7F-G). Overall, these *in vivo* experiments further highlight a role for Serpina3n in BBB dysfunction.

**Figure 7.**
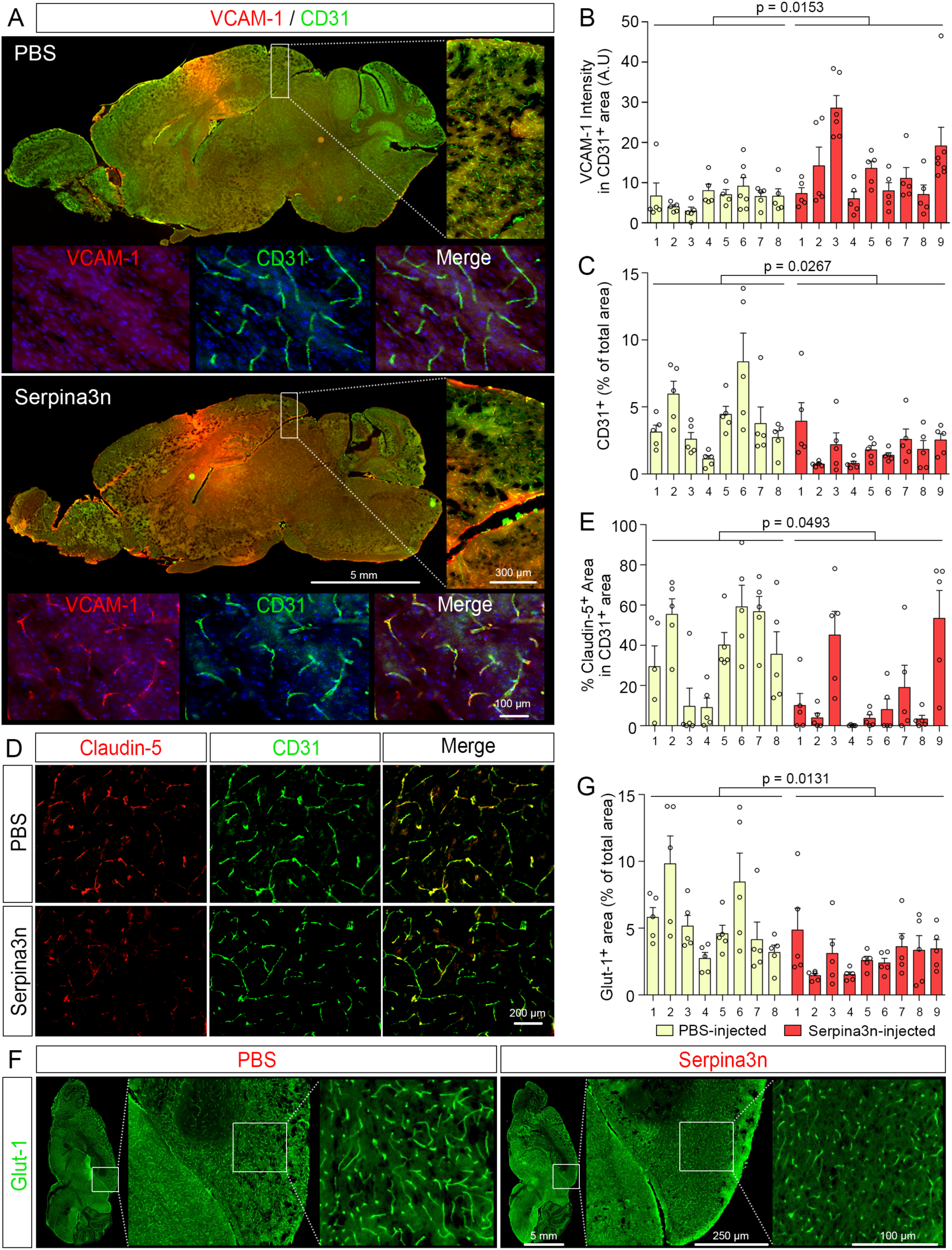
Exogenously administered Serpina3n causes BBB dysfunction *in vivo*. N=8 mice received an ICV injection of PBS and N=9 mice received an ICV injection of Serpina3n, and the entire cohort was euthanized 4 days after injections. For all quantifications, 5-7 regions in each mouse were randomly imaged at a distance of 3 mm away from the needle track to minimize contributions from sterile inflammation. (A) Representative sagittal brain section highlighting increased VCAM-1 expression after delivery of Serpina3n. The inset shows that VCAM-1 is localized exclusively to CD31^+^ blood vessels. (B) Quantification of VCAM-1 intensity within total CD31^+^ area. Each data point represents an individual image, and data are graphed as mean ± SEM for each biological replicate. Statistical significance was calculated by a student’s unpaired t-test. (C) Quantification of vessel coverage by normalizing CD31^+^ area to total tissue area. Each data point represents an individual image, and data are graphed as mean ± SEM for each biological replicate. Statistical significance was calculated by a student’s unpaired t-test. (D) Representative images of CD31 and claudin-5 expression in a sagittal brain section. (E) Quantification of claudin-5 coverage within CD31^+^ vessel area. Each data point represents an individual image, and data are graphed as mean ± SEM for each biological replicate. Statistical significance was calculated by a student’s unpaired t-test. (F) Representative images of Glut-1 expression in a sagittal brain section. (G) Normalized Glut-1^+^ area. Each data point represents an individual image, and data are graphed as mean ± SEM for each biological replicate. Statistical significance was calculated by a student’s unpaired t-test.

## Discussion

Recently, it has been proposed that proinflammatory conditions generate subsets of inflammatory reactive astrocytes that may be active participants in neurodegenerative disease progression. Inflammation in brain endothelial cells is also a common feature of age-related neurodegenerative diseases. However, a detailed understanding of the molecular basis and interactions between subtypes of reactive astrocytes and brain endothelial cells has remained largely unknown. Progress in this area has been limited by the heterogeneity of reactive astrocytes *in vivo*, as well as the absence of effective *in vitro* systems that permit long-term quiescent cocultures of astrocytes and brain endothelial cells that can be perturbed in a controlled manner. Indeed, *in vitro* cultures of astrocytes and brain endothelial cells have historically relied on serum, an inducer of astrocyte reactivity^74, 78^, which prevented defined cause-and-effect studies of astrocyte/endothelial crosstalk. Herein, we focused on a simplified human BBB model generated from iPSCs, which offers a scalable system that also permits tight control over experimental conditions. The differentiation of iPSCs to BMEC-like cells and astrocytes has been iterated over the past decade, and a recent step forward that enabled this current study was our prior establishment of a serum-free protocol yielding BMEC-like cells with representative BBB characteristics. This allowed us to use a variety molecular tools and analyses to understand BBB alterations to astrocyte reactivity.

Previous studies have shown that iPSC-derived BMEC-like cells do not express VCAM-1 on the cell surface after TNFα stimulation^40^, in contrast to primary BMECs that readily upregulate VCAM-1^79^. After validating that we could convert iPSC-derived astrocytes to a GBP2^+^ inflammatory reactive state, we studied whether astrocytes could influence BMEC responses to TNFα. We ultimately determined that active astrocyte/endothelial crosstalk was necessary for TNFα to disrupt passive BBB function and yield upregulation of VCAM-1 on the BMEC-like cells. These findings suggested BBB inflammation is not fully cell autonomous, a notion supported by other studies^80, 81^.

Activation of STAT3 signaling has been suggested as a central axis for a subpopulation of reactive astrocytes in CNS disorders^60, 61, 82, 83^. For example, in mouse models, STAT3 signaling in reactive astrocytes has been tied motor outcomes in stroke^84^, and plaque burden and memory deficiencies in AD^60^. In human patients, STAT3 activation in astrocytes regulates cancer metastasis to brain^85^. The severity and progression of many brain diseases are also linked human neurovascular dysfunction^86, 87^, which suggested links between STAT3 activation in astrocytes and BBB dysfunction. Our *in vitro* data showing rescue of BBB properties through inhibition of STAT3 and its downstream targets in astrocytes, as well as our human tissue assessments, support this link. Our merged investigations into STAT3-dependent astrocyte reactivity and BBB dysfunction further allowed us to identify α1ACT/Serpina3n as a novel BBB-damaging factor. Historically, Serpina3n and α1ACT have been associated with astrocytes in mouse models of AD^88^ and postmortem human tissue^89^, respectively, and these proteins are believed to play a role in amyloid beta aggregation^90^. More recent studies have linked Serpina3n to factors associated with BBB dysfunction, without providing direct links between Serpina3n and BBB dysfunction. For example, the APOE4 genotype yields increased expression of Serpina3n in humanized APOE mice relative to APOE3 and APOE2^91^, and in separate work, APOE4 humanized mice exhibited increased BBB disruption relative to APOE3 and APOE2 mice through a cyclophilin A-dependent inflammatory cascade^92^. APOE4 has also recently been identified as a driver of BBB dysfunction and cognitive impairment in human subjects independent of amyloid and tau pathology,^93^ which could potentially be linked to α1ACT. Aging and intraperitoneal injection of lipopolysaccharide, which yield loss of BBB function, also produce an increase in Serpina3n expression in astrocytes across many different brain regions^18, 77, 94^. Hence, we speculate that astrocyte-derived Serpina3n/α1ACT could be an unrecognized driver of BBB dysfunction in many diseases. Indeed, α1ACT was recently identified as a potential biomarker for progressive for multiple sclerosis due to its elevated levels in cerebrospinal fluid^95^, which could be connected to increased VCAM-1 expression at the BBB to license immune cell entry into the brain.

We further suggest that establishing more robust connections between reactive astrocyte subsets and BBB dysfunction across diverse animal models and human tissue cohorts will be necessary to continuously shape our understanding for how astrocyte-BBB crosstalk contributes to human neurodegeneration. For example, reactive astrocytes were recently shown to support vascular repair and remodeling after stroke, with worsened outcomes if reactive astrocytes were ablated^96^. This animal model of stroke utilized photoablation rather than artery occlusion, which is the stroke model in which Serpina3n was identified as a reactive astrocyte marker^74^, and *Serpina3n* was not highlighted in the photoablation datasets. Hence, the specificity of the injury may be key for induction of STAT3 signaling and Serpina3n secretion by the resultant reactive astrocytes. We note that the reactive astrocyte state characterized in our study aligns with the “STAT3-dependent reactivity” state identified by Kun et al, which is defined by an acute phase inflammatory response to IL-1*α*, TNF*α*, and C1q leading to upregulation of similar genes^72^. Based on the results, there may also be specific reactive astrocyte subtypes within the STAT3-dependent group that could be further interrogated for a role in BBB dysfunction by mechanisms that are independent of Serpina3n/α1ACT. These possibilities are well-suited for future exploration using the comprehensive framework established in our study.

## Material and Methods

### iPSC maintenance

The CC3 iPSC line was used for all experiments unless otherwise stated^97^. Undifferentiated iPSCs were maintained in E8 medium on 6-well plates coated with growth factor reduced Matrigel (Corning). iPSCs were passaged with Versene (Thermo Fisher Scientific) upon reaching 60-80% confluency.

### iPSC differentiation to astrocytes

To reduce variability in astrocyte generation, manually isolated neural progenitors were used as the starting population. As shown in Figure 2A, iPSCs were dissociated with Versene and seeded into 6-well low-attachment plates to form embryoid bodies (EBs) at a density of 4×10^5^ cells/well in E6 medium supplemented with 100x N2 (Invitrogen; E6N2) and 10 μM Y27632 (Tocris). Neural differentiation in the EBs was achieved by dual inhibition of SMAD signaling for 7 days^98^. EBs were then seeded onto growth factor reduced Matrigel-coated plates in E6N2 containing 1 μg/ml laminin (Sigma Aldrich). After another 7 days (Day 0), neural progenitor cells were manually isolated from rosettes and expanded as neurospheres in a suspension culture for 7 days in E6N2 supplemented with 50x B27 lacking retinoic acid (Invitrogen) and 20 ng/ml FGF2 (Peprotech). Then, the neurospheres were dissociated into single cells with TrypLE™ (Thermo Fisher Scientific) and seeded onto growth factor reduced Matrigel-coated 6-well plates in E6N2 supplemented with 50x B27 lacking retinoic acid, 10 ng/ml BMP4 (Peprotech) and 20 ng/ml FGF2 for directed astroglial differentiation^99^. Medium changes were performed every 48 hours, and cells were passaged upon reaching approximately 80% confluency. Astrocytes were seeded for coculture at a ratio of 1 well of a 6-well plate to 6 wells of a 12-well plate and maintained in E6 medium containing 10 ng/ml CNTF (Peprotech) and 10 ng/ml EGF (Peprotech) until initiation of coculture.

### iPSC differentiation to BMEC-like cells

Differentiations were carried out as previously described^35^. iPSCs were dissociated with Accutase and seeded onto growth factor reduced Matrigel-coated plates in E8 medium containing 10 μM Y27632 at a density of 15,000 cells/cm^2^. Differentiation was initiated 24 hours after seeding by changing to E6 medium, with daily medium changes for 4 days. Next, cells were expanded with serum-free basal endothelial cell medium (EC medium) supplemented with 50x B27 (Thermo Fisher Scientific), GlutaMAX™ (Thermo Fisher Scientific), 10 μM retinoic acid (Sigma Aldrich), and 20 μg/ml FGF2 for 2 days without media change. Following this treatment, cells were collected by a 20-minute incubation in Accutase and seeded onto Transwell filters (1.1 cm^2^ polyethylene terephthalate membranes with 0.4 μm pores; Fisher Scientific) coated with a mixture of 400 μg/ml collagen IV (Sigma Aldrich) and 100 μg/ml fibronectin (Sigma Aldrich). The following day, cells were switched to EC medium lacking FGF2 and RA. If coculture was being initiated, filters were transferred to 12-well plates containing astrocytes and the same medium was utilized. Starting at this time, transendothelial electrical resistance (TEER) was measured using STX2 chopstick electrodes and an EVOM2 voltameter (World Precision Instruments) and approximately every 24 hours thereafter.

### Cytokine treatments on astrocytes and BMEC-like cells

To prepare conditioned medium, astrocyte cultures at 80% confluence in 6-well plates (in E6 medium containing BMP4 and FGF2) were treated with vehicle control or IL-1a (5 ng/ml; Peprotech), TNFα (10 ng/ml; Peprotech), and C1a (500 ng/ml; MyBioSource) for 72 hours. After three washes with DPBS, cells were placed in minimal conditioning medium containing phenol-red-free DMEM/F12 (Thermo Fisher Scientific) and Glutamax (Gibco). Spent media were collected after 3 days, with cell debris removed by centrifugation at 3000 RPM, 4°C for 5 minutes. The supernatants were concentrated in centrifugal concentrators (Millipore Sigma) with a size cutoff filter of 3 kDa. Protein concentration was determined by a BCA assay (Thermo Fisher Scientific), and the solutions were added to BMEC-like cells at 100 μg/ml. For direct treatment of cytokines on either BMECs in monoculture or coculture with astrocytes, a full media change with vehicle control or cytokine (5 ng/ml IL-1a, 10 ng/ml TNFα, 500 ng/ml of C1q, or 10 ng/ml of IFNγ) was performed a day after seeding of BMEC-like cells onto Transwell filters as described above.

### Generation of adeno-associated virus (AAV)

Plasmid pAdDeltaF6 was a gift from James Wilson (Addgene plasmid #112867). pUCmini-iCAP-PHP.B was a gift from Viviana Gradinaru (Addgene plasmid #103002). Plasmid GFAP(short)-shRNAmiR was constructed by VectorBuilder Inc. The shRNAmiR sequences in transfer vectors were incorporated into AAV particles using pUCmini-iCAP-PHP.B and pAdDeltaF6. Virus was produced by transfecting all plasmids at a 1:1:1 molar ratio (30 μg total DNA) into HEK293T cells with Lipofectamine3000 (Invitrogen). Viral particles were collected at 72 hours by filtering spent medium through a 0.45 μm cellulose acetate filter (Millipore Sigma), then concentrating the supernatant with AAVpro® concentrator (TaKaRa).

### Generation of lentivirus

Plasmid EF.STAT3DN.UBC.GFP, which contains the cDNA encoding for the dominant-negative form of STAT3 (STAT3DN), was a gift from Linzhao Cheng (Addgene plasmid #24984). Plasmid pMK1334 for lentiviral delivery of sgRNA was a gift from Martin Kampmann (Addgene plasmid #127965). Sequences in transfer vectors were incorporated into lentivirus using the lentiviral packaging system (pMD2.G and psPAX2), which was a gift from Didier Trono (Addgene plasmid #12259 and #12260) in HEK293T cells using Lipofectamine 3000 (Invitrogen). After 48 and 72 hours, spent medium was collected and filtered through a 0.45 μm cellulose acetate filter before concentrating the supernatant using Lenti-X™ concentrator (TaKaRa).

### RNA sequencing and data analysis

All samples were collected at the day 5 time point indicated in Figure 2J, regardless of monoculture or coculture. Total RNA was isolated using TRIzol (Sigma Aldrich). RNA was further purified with a Qiagen RNeasy Plus mini kit (Qiagen) and submitted to the Vanderbilt Technologies for Advanced Genomics (VANTAGE) core for sequencing using an Illumina NovaSeq6000. Samples were prepared for sequencing using the TruSeq RNA sample prep kit (Illumina) to prepare cDNA libraries after ribosome depletion. Raw sequencing reads were obtained for the paired-end samples. FASTQ reads were mapped to the human genome (hg19) by HISAT2 (2.2.0). EdgeR (3.30.3) and limma^100^ software packages were used to measure differential gene expression with genes that achieved a count per million mapped reads (CPM). Any genes not considered to be detected (CPM less than 4) were removed. Adjusted p-values (<0.05) were utilized for functional enrichment analysis with the WEB-based Gene SeT AnaLysis Toolkit (WebGestalt)^101^. For interactome prediction between astrocytic secretome and endothelial receptome, the STRING database^102^ was used with combined edge confidence scores of >0.7 of the human interactome version 11.0b. Network figures were created using Origene2021 where nodes refer to genes through identified protein-protein interaction networks. For an unbiased analysis of potential differentially regulated pathways in astrocytes, differentially expressed genes (FDR<0.1) and corresponding expression values were uploaded into the Ingenuity Pathway Analysis system (http://www.ingenuity.com).

### qPCR analysis

Total RNA was extracted with TRIzol and converted to cDNA using a Superscript III First-Strand kit (Invitrogen). qPCR was performed on a BioRad CFX96 Thermocycler with TaqMan primers against *VEGFA*, *VCAM1*, and *GAPDH* (Thermo Fisher Scientific). Samples were analyzed by normalizing expression levels to *GAPDH* and relative quantification was performed using the standard 2-ΔΔC_t_ method.

### STAT3 signaling inhibition experiments

For experiments involving overexpression of dominant-negative STAT3 _Y705F_, freshly plated iPSC-derived astrocytes (10^6^ cells per 6-well plate) were incubated for three days with lentiviral particles expressing dominant-negative STAT3_Y705F_ in E6 medium containing 10 ng/ml BMP4 and 10 ng/ml EGF. Astrocytes were expanded and maintained by changing medium every 48 hours until initiation of coculture with BMEC-like cells. Empty viral particles were used as a control. Expression of EGFP in astrocytes was confirmed on an epifluorescent microscope before proceeding with experiments.

### CRISPR interference (CRISPRi) experiments

WTC11 iPSCs with stably integrated CRISPRi machinery and dox-inducible NFIA-SOX9^72^ were dissociated to a single-cell suspension with Accutase and then replated at 7,500 cells/cm^2^ in Matrigel-coated 10-cm or 15-cm dishes with DMEM/F12 basal medium supplemented with 50x B27, 100x N2, and 20 μg/ml FGF2. The next day, media was changed to ScienCell Astrocyte Media (ScienCell Research Laboratories) containing 2 μg/ml doxycycline (Millipore Sigma) to initiate astrocyte differentiation. A full media exchange was conducted every other day, with doxycycline maintained at 2 μg/ml throughout the differentiation process. In 5-6 days, after confluency was reached, the cultures were dissociated and split 1:8. Expansion of the cultures with the same split ratio was continued until day 20 of differentiation. The final CRISPRi-astrocytes were then characterized as described previously^72^. Prior to coculture with BMEC-like cells, CRISPRi-astrocytes were incubated with lentiviral particles for 3 days and then passaged once with media exchanges every other day. The following sgRNAs were cloned into the pMK1334 vector before lentiviral production:

*JAK3* (GACCTAGCGGGCAGGGACCC)
*DSP* (GAACTCGGACGGCTACTGGT)
*SOD2* (GGGCGCAGGAGCGGCACTCG)
*CCL2* (GAGCAGCCAGAGGAACCGAG)
*CXCL6* (GGTGGAGGCGCGGAGACTGG)
*CXCL8* (GAGAGCCAGGAAGAAACCAC)
*CXCL10* (GCTGGAGGTTCCTCTGCTGT)
*C3* (GTGCAGGGTCAGAGGGACAG)
*SERPINA3* (GAAAAGGCACAGGGAATCAG)
*SLC43A2* (GAGATCAGCGCCCAGACCTA)
*TNFAIP2* (GAATGGACGTGTGGAAGCGG)
*VEGFA* (GGGGCAGCCGGGTAGCTCGG)

### Human postmortem brain tissue

Brain tissue collection procedures were approved by the Institutional Review Board at Vanderbilt University Medical Center, and written consent for brain donation was obtained from patients or their surrogate decision-makers. A diagnostic postmortem evaluation was performed to confirm the presence of Alzheimer’s disease or cerebral amyloid angiopathy following the National Alzheimer’s Coordinating Center (NACC) Neuropathology Data Form. The tissue was flash-frozen in liquid nitrogen at the time of donation. De-identified brain tissue from a patient with unspecified non-surgical convulsive seizures was obtained via Vanderbilt University Medical Center Cooperative Human Tissue Network.

### Slice culture preparation and use

All animal procedures were performed in accordance with a protocol approved by the Institutional Animal Care and Use Committee at Vanderbilt University. P6 C57BL/6J mice (both male and female pups) were sacrificed by cervical dislocation. Brains were removed, mounted, and cut into 300 μm coronal slices. Transverse slices of 300 μm were made on a vibrating blade microtome (Leica) in cold Hank’s Balanced Salt Solution supplemented with 100 U/ml penicillin/streptomycin, 10 mM HEPES, and 4.5 mM sodium bicarbonate. Each slice was transferred to a membrane insert (PICMORG50, Millipore Sigma) in a 35 mm culture dish with 1.2 ml of DMEM supplemented with 3.5 g/L D-glucose, 0.2 mM Glutamax, 10 mM HEPES, 25 U/ml penicillin/streptomycin, 50x B27 Plus, and 0.1 mM D-Luciferin sodium salt (Tocris). For knockdown of Serpina3n, slice cultures were transduced with AAV particles at the onset of culture. AAV particles harbored an shRNA sequence targeting *Serpina3n* (AACGGGTAGTGCCCTGTTTATT) or the original shRNA sequence targeting LacZ as a negative control (TGTCGGCTTACGGCGGTGATTT). Three days later, the slice cultures were stimulated for another three days with 10 ng/ml TNFα prior to analysis. For Serpina3n dose-response assays, recombinant mouse Serpina3n (R&D Systems), at a concentration of 50 or 100 ng/ml, was added at the onset of culture and incubated for four days. For all experiments, slice chambers were sealed with a coverslip and maintained in a humidified incubator.

### Serpina3n delivery to brain via intravecerebroventricular injection

Four-month-old wild-type C57BL/6J mice males (Jackson Laboratories) were randomly assigned to the control (PBS) and Serpina3n-injected groups (n=8-9 per group). Under isoflurane anesthesia, mice received a stereotactic injection directly into the ventricle at coordinates of AP = -0.3 mm, ML = -1 mm, and DV = -3 mm. After injections, mice were immediately returned to prewarmed cages. The first group received an injection of 4 μl of sterile saline solution and the second group received an injection of 10 ng/μl Serpina3n solution (40 ng in 4 total μl). Four days after surgery, mice were perfused with PBS and sacrificed for brain extraction and histology.

### Immunostaining

Cryosectioned tissues, slice cultures, and cultured cells were fixed with 4% paraformaldehyde and processed for immunofluorescence staining. Samples were permeabilized with 0.3% Triton X-100 in PBS, blocked with 10% goat serum in PBS, and incubated at 4°C overnight with the following primary antibodies: mouse anti-C3 (1:200 dilution, 846302, Biolegend), rabbit anti-CD31 (1:200 dilution, RB-10333, Thermo), mouse anti-CD44 (1:1000 dilution, AB6124, Abcam), anti-Claudin-5-Alexa Fluor 488 (1:1000 dilution, 352488, Invitrogen), mouse anti-GBP2 (1:100 dilution, LS-B12172-50, LSBio), rabbit anti-GFAP (1:300 dilution, Z0334, Dako), mouse anti-GFAP (1:300 dilution, MAB360, Millipore), chicken anti-GFAP (1:300 dilution, SKU: GFAP, Aves), goat anti-GFP (1:1000 dilution, 600-141-215, Rockland), mouse anti-Glut1 (1:1000 dilution, FAB1418G, R&D systems), mouse anti-occludin (1:500 dilution, 33-1500, Thermo Fisher Scientific), rabbit anti-pSTAT3(y705) (1:300 dilution, 9145S, Cell Signaling), goat anti-Serpina3n (1:200 dilution, AF4709-SP, Abcam), rabbit anti-VCAM-1 (1:200 dilution, ab134047, Abcam), mouse anti-VCAM-1 (1:200 dilution, sc-13160, Santa Cruz), goat anti-VE-cadherin (1:300 dilution, AF938, R&D Systems). The following day, after washing with PBS, for unconjugated antibodies, immunostaining was completed by a 1-hour room temperature incubation with secondary antibody (donkey anti-rabbit, anti-mouse, anti-chicken, or anti-goat Alexa Fluor 488, Alexa Fluor 546, or Alexa Fluor 647; 1:1,000 dilution; Thermo Fisher Scientific). Tissue sections were mounted with the anti-fade Fluoromount-G medium containing 1,40,6-diamidino-2-phenylindole dihydrochloride (DAPI; Southern Biotechnology), while cultured cells were labeled with DAPI (Thermo Fisher Scientific) diluted in PBS for 10 minutes. Images were acquired with a Zeiss LSM700 T-PMT confocal microscope or a Leica DMi8 epifluorescence microscope. In select cases, images were analyzed using customized scripts for the macro function of ImageJ software (1.53c).

### Western blot analysis

BMEC-like cells were lysed with RIPA buffer (Thermo Fisher Scientific) supplemented with protease inhibitor (Sigma Aldrich). Lysed protein was separated on 12% SDS-PAGE gels and transferred onto nitrocellulose membranes. Blots processed with blocking solution were incubated with antibodies against VCAM-1 (ab134047, Abcam, 1:1000 dilution), E-Selectin (AF724, R&D systems, 1:1000 dilution), and GAPDH (D16H11, Cell Signaling, 1:5000 dilution). The membranes were then washed and incubated with LI-COR fluorophore-conjugated second antibodies (926-68024, 926-32214, 926-68022, 926-32210, and 926-32211). Afterward, western blots were visualized using an Odyssey® Fc (LI-COR).

### Enzyme-linked immunosorbent assay detection of Serpina3n

Slice cultures were treated with AAV particles for 3 days, followed by TNFα or vehicle control for 3 days, then spent medium was collected. A Serpina3n-specific sandwich ELISA kit (Aviva Biosciences) was used according to the manufacturer’s recommendation. Briefly, spent media were allowed to react with a capture anti-Serpina3n antibody on the surface of 96-well ELISA plates for 2 hours at room temperature. A second anti-Serpina3n antibody was used to detect immobilized Serpina3n, followed by incubation with an HRP-conjugated antibody and detection at 450 nm. Quantification was achieved using a standard curve.

## Author Contributions

HK and ESL conceived the study and designed all experiments. HK, AGS, AS, SMS, and EHN carried out experiments. KL and MK provided key reagents and input for the CRISPRi experiments. JP provided assistance with the bioinformatics analyses. SK and DGM provided slice cultures and input on their use. MSS provided human tissue samples and assisted with histological analyses.

## Acknowledgments

Funding was provided by Chan Zuckerberg Initiative Ben Barres Early Career Acceleration Awards to ESL and MK, as well as grants from the National Institutes of Health (R01 GM117650 to DGM, F30 AG066418 to KL, R21 AG070859 and K76 AG060001 to MSS). SK was supported by a Vanderbilt International Scholarship. EHN was supported by a National Science Foundation Graduate Research Fellowship. Support for RNA sequencing was provided by the Vanderbilt Technologies for Advanced Genomics core facility, which is supported in part by a Clinical and Translational Science Award (5UL1 RR024975), the Vanderbilt Ingram Cancer Center (P30 CA68485), the Vanderbilt Vision Center (P30 EY08126), a CTSA award from the National Center for Advancing Translational Sciences (UL1 TR002243), and the National Center for Research Resources (G20 RR030956). Pilot funding was also provided under CTSA award UL1 TR002243. The contents of this manuscript are solely the responsibility of the authors and do not necessarily represent official views of the National Institutes of Health.

## Competing interest statement

MK serves on the Scientific Advisory Board of Engine Biosciences, Casma Therapeutics, and Cajal Neuroscience, and is an advisor to Modulo Bio and Recursion Therapeutics. None of the other authors declare competing interests.

## Data availability statement

RNA sequencing data have been uploaded to ArrayExpress under the accession number E-MTAB-11468.

## References

1. Hawkins, B.T. & Davis, T.P. The blood-brain barrier/neurovascular unit in health and disease. Pharmacol Rev 57, 173–185 (2005).

2. Stanimirovic, D.B. & Friedman, A. Pathophysiology of the neurovascular unit: disease cause or consequence? J Cereb Blood Flow Metab 32, 1207–1221 (2012).

3. Najjar, S., Pearlman, D.M., Devinsky, O., Najjar, A. & Zagzag, D. Neurovascular unit dysfunction with blood-brain barrier hyperpermeability contributes to major depressive disorder: a review of clinical and experimental evidence. J Neuroinflammation 10, 142 (2013).

4. Sweeney, M.D., Ayyadurai, S. & Zlokovic, B.V. Pericytes of the neurovascular unit: key functions and signaling pathways. Nature Neuroscience 19, 771–783 (2016).

5. Pulido, R.S., et al. Neuronal Activity Regulates Blood-Brain Barrier Efflux Transport through Endothelial Circadian Genes. Neuron 108, 937–952.e937 (2020).

6. Abbott, N.J., Rönnbäck, L. & Hansson, E. Astrocyte-endothelial interactions at the blood-brain barrier. Nat Rev Neurosci 7, 41–53 (2006).

7. Janzer, R.C. & Raff, M.C. Astrocytes Induce Blood-Brain-Barrier Properties in Endothelial-Cells. Nature 325, 253–257 (1987).

8. Hayashi, Y., et al. Induction of various blood-brain barrier properties in non-neural endothelial cells by close apposition to co-cultured astrocytes. Glia 19, 13–26 (1997).

9. Alvarez, J.I., et al. The Hedgehog pathway promotes blood-brain barrier integrity and CNS immune quiescence. Science 334, 1727–1731 (2011).

10. Wang, H., et al. Inactivation of Hedgehog signal transduction in adult astrocytes results in region-specific blood-brain barrier defects. Proc Natl Acad Sci U S A 118 (2021).

11. Wosik, K., et al. Angiotensin II controls occludin function and is required for blood brain barrier maintenance: relevance to multiple sclerosis. J Neurosci 27, 9032–9042 (2007).

12. Mi, H., Haeberle, H. & Barres, B.A. Induction of astrocyte differentiation by endothelial cells. J Neurosci 21, 1538–1547 (2001).

13. Kubotera, H., et al. Astrocytic endfeet re-cover blood vessels after removal by laser ablation. Sci Rep 9, 1263 (2019).

14. Heithoff, B.P., et al. Astrocytes are necessary for blood-brain barrier maintenance in the adult mouse brain. Glia 69, 436–472 (2021).

15. Hartmann, K., et al. Complement 3(+)-astrocytes are highly abundant in prion diseases, but their abolishment led to an accelerated disease course and early dysregulation of microglia. Acta Neuropathol Commun 7, 83 (2019).

16. Ugalde, C.L., et al. Markers of A1 astrocytes stratify to molecular sub-types in sporadic Creutzfeldt-Jakob disease brain. Brain Commun 2, fcaa029 (2020).

17. Liddelow, S.A., et al. Neurotoxic reactive astrocytes are induced by activated microglia. Nature 541, 481–487 (2017).

18. Guttenplan, K.A., et al. Knockout of reactive astrocyte activating factors slows disease progression in an ALS mouse model. Nat Commun 11, 3753 (2020).

19. Yun, S.P., et al. Block of A1 astrocyte conversion by microglia is neuroprotective in models of Parkinson’s disease. Nat Med 24, 931–938 (2018).

20. Miyamoto, N., et al. The effects of A1/A2 astrocytes on oligodendrocyte linage cells against white matter injury under prolonged cerebral hypoperfusion. Glia 68, 1910–1924 (2020).

21. Chapouly, C., et al. Astrocytic TYMP and VEGFA drive blood-brain barrier opening in inflammatory central nervous system lesions. Brain 138, 1548–1567 (2015).

22. Chang, C.Y., et al. Disruption of in vitro endothelial barrier integrity by Japanese encephalitis virus-Infected astrocytes. Glia 63, 1915–1932 (2015).

23. Deng, Z., et al. Astrocyte-derived VEGF increases cerebral microvascular permeability under high salt conditions. Aging (Albany NY) 12, 11781–11793 (2020).

24. Persidsky, Y., et al. Microglial and astrocyte chemokines regulate monocyte migration through the blood-brain barrier in human immunodeficiency virus-1 encephalitis. Am J Pathol 155, 1599–1611 (1999).

25. Hudson, L.C., Bragg, D.C., Tompkins, M.B. & Meeker, R.B. Astrocytes and microglia differentially regulate trafficking of lymphocyte subsets across brain endothelial cells. Brain Res 1058, 148–160 (2005).

26. Liu, C.Y., Yang, Y., Ju, W.N., Wang, X. & Zhang, H.L. Emerging Roles of Astrocytes in Neuro-Vascular Unit and the Tripartite Synapse With Emphasis on Reactive Gliosis in the Context of Alzheimer’s Disease. Front Cell Neurosci 12, 193 (2018).

27. Cabezas, R., et al. Astrocytic modulation of blood brain barrier: perspectives on Parkinson’s disease. Front Cell Neurosci 8, 211 (2014).

28. Myer, D.J., Gurkoff, G.G., Lee, S.M., Hovda, D.A. & Sofroniew, M.V. Essential protective roles of reactive astrocytes in traumatic brain injury. Brain 129, 2761–2772 (2006).

29. Linnerbauer, M. & Rothhammer, V. Protective Functions of Reactive Astrocytes Following Central Nervous System Insult. Front Immunol 11, 573256 (2020).

30. Bush, T.G., et al. Leukocyte infiltration, neuronal degeneration, and neurite outgrowth after ablation of scar-forming, reactive astrocytes in adult transgenic mice. Neuron 23, 297–308 (1999).

31. Bosworth, A.M., Faley, S.L., Bellan, L.M. & Lippmann, E.S. Modeling Neurovascular Disorders and Therapeutic Outcomes with Human-Induced Pluripotent Stem Cells. Front Bioeng Biotechnol 5, 87 (2017).

32. Perriot, S., et al. Human Induced Pluripotent Stem Cell-Derived Astrocytes Are Differentially Activated by Multiple Sclerosis-Associated Cytokines. Stem Cell Reports 11, 1199–1210 (2018).

33. Barbar, L., et al. CD49f Is a Novel Marker of Functional and Reactive Human iPSC-Derived Astrocytes. Neuron 107, 436–453.e412 (2020).

34. Lippmann, E.S., Azarin, S.M., Palecek, S.P. & Shusta, E.V. Commentary on human pluripotent stem cell-based blood-brain barrier models. Fluids Barriers CNS 17, 64 (2020).

35. Neal, E.H., et al. A Simplified, Fully Defined Differentiation Scheme for Producing Blood-Brain Barrier Endothelial Cells from Human iPSCs. Stem Cell Reports 12, 1380–1388 (2019).

36. Neal, E.H., et al. Influence of basal media composition on barrier fidelity within human pluripotent stem cell-derived blood-brain barrier models. J Neurochem 159, 980–991 (2021).

37. Cortes-Canteli, M. & Iadecola, C. Alzheimer’s Disease and Vascular Aging: JACC Focus Seminar. J Am Coll Cardiol 75, 942–951 (2020).

38. Sweeney, M.D., Sagare, A.P. & Zlokovic, B.V. Blood-brain barrier breakdown in Alzheimer disease and other neurodegenerative disorders. Nat Rev Neurol 14, 133–150 (2018).

39. Martins Gomes, S.F., et al. Induced Pluripotent Stem Cell-Derived Brain Endothelial Cells as a Cellular Model to Study. Front Microbiol 10, 1181 (2019).

40. Nishihara, H., et al. Advancing human induced pluripotent stem cell-derived blood-brain barrier models for studying immune cell interactions. FASEB J 34, 16693–16715 (2020).

41. Ghimire, K., et al. CD47 Promotes Age-Associated Deterioration in Angiogenesis, Blood Flow and Glucose Homeostasis. Cells 9 (2020).

42. Widagdo, J., Fang, H., Jang, S.E. & Anggono, V. PACSIN1 regulates the dynamics of AMPA receptor trafficking. Sci Rep 6, 31070 (2016).

43. Barbera, S., et al. The small GTPase Rab5c is a key regulator of trafficking of the CD93/Multimerin-2/β1 integrin complex in endothelial cell adhesion and migration. Cell Commun Signal 17, 55 (2019).

44. Lu, Q., et al. Angiogenic Factor AGGF1 Activates Autophagy with an Essential Role in Therapeutic Angiogenesis for Heart Disease. PLoS Biol 14, e1002529 (2016).

45. Gatina, D.Z., et al. Proangiogenic Effect of 2A-Peptide Based Multicistronic Recombinant Constructs Encoding VEGF and FGF2 Growth Factors. Int J Mol Sci 22 (2021).

46. Coffelt, S.B., et al. Angiopoietin-2 regulates gene expression in TIE2-expressing monocytes and augments their inherent proangiogenic functions. Cancer Res 70, 5270–5280 (2010).

47. Sloan, S.A., et al. Human Astrocyte Maturation Captured in 3D Cerebral Cortical Spheroids Derived from Pluripotent Stem Cells. Neuron 95, 779–790.e776 (2017).

48. Del-Aguila, J.L., et al. A single-nuclei RNA sequencing study of Mendelian and sporadic AD in the human brain. Alzheimers Res Ther 11, 71 (2019).

49. Ulbrich, H., Eriksson, E.E. & Lindbom, L. Leukocyte and endothelial cell adhesion molecules as targets for therapeutic interventions in inflammatory disease. Trends Pharmacol Sci 24, 640–647 (2003).

50. Yousef, H., et al. Aged blood impairs hippocampal neural precursor activity and activates microglia via brain endothelial cell VCAM1. Nat Med 25, 988–1000 (2019).

51. Weksler, B.B., et al. Blood-brain barrier-specific properties of a human adult brain endothelial cell line. FASEB J 19, 1872–1874 (2005).

52. Chen, G., et al. Comprehensive Identification and Characterization of Human Secretome Based on Integrative Proteomic and Transcriptomic Data. Front Cell Dev Biol 7, 299 (2019).

53. Jin, W. Role of JAK/STAT3 Signaling in the Regulation of Metastasis, the Transition of Cancer Stem Cells, and Chemoresistance of Cancer by Epithelial-Mesenchymal Transition. Cells 9 (2020).

54. Yeruva, S., Ramadori, G. & Raddatz, D. NF-kappaB-dependent synergistic regulation of CXCL10 gene expression by IL-1beta and IFN-gamma in human intestinal epithelial cell lines. Int J Colorectal Dis 23, 305–317 (2008).

55. Kanda, N., Shimizu, T., Tada, Y. & Watanabe, S. IL-18 enhances IFN-gamma-induced production of CXCL9, CXCL10, and CXCL11 in human keratinocytes. Eur J Immunol 37, 338–350 (2007).

56. Liu, X., et al. Stat3 inhibition attenuates mechanical allodynia through transcriptional regulation of chemokine expression in spinal astrocytes. PLoS One 8, e75804 (2013).

57. Gu, F.M., et al. IL-17 induces AKT-dependent IL-6/JAK2/STAT3 activation and tumor progression in hepatocellular carcinoma. Mol Cancer 10, 150 (2011).

58. Kulesza, D.W., et al. Search for novel STAT3-dependent genes reveals SERPINA3 as a new STAT3 target that regulates invasion of human melanoma cells. Lab Invest 99, 1607–1621 (2019).

59. Szklarczyk, D., et al. The STRING database in 2011: functional interaction networks of proteins, globally integrated and scored. Nucleic Acids Res 39, D561–568 (2011).

60. Reichenbach, N., et al. Inhibition of Stat3-mediated astrogliosis ameliorates pathology in an Alzheimer’s disease model. EMBO Mol Med 11 (2019).

61. Ben Haim, L., et al. The JAK/STAT3 pathway is a common inducer of astrocyte reactivity in Alzheimer’s and Huntington’s diseases. J Neurosci 35, 2817–2829 (2015).

62. Choi, M., Kim, H., Yang, E.J. & Kim, H.S. Inhibition of STAT3 phosphorylation attenuates impairments in learning and memory in 5XFAD mice, an animal model of Alzheimer’s disease. J Pharmacol Sci 143, 290–299 (2020).

63. Wen, Z., Zhong, Z. & Darnell, J.E. Maximal activation of transcription by Stat1 and Stat3 requires both tyrosine and serine phosphorylation. Cell 82, 241–250 (1995).

64. Argaw, A.T., et al. Astrocyte-derived VEGF-A drives blood-brain barrier disruption in CNS inflammatory disease. J Clin Invest 122, 2454–2468 (2012).

65. Kohli, K., Pillarisetty, V.G. & Kim, T.S. Key chemokines direct migration of immune cells in solid tumors. Cancer Gene Ther (2021).

66. Deshmane, S.L., Kremlev, S., Amini, S. & Sawaya, B.E. Monocyte chemoattractant protein-1 (MCP-1): an overview. J Interferon Cytokine Res 29, 313–326 (2009).

67. Lu, H., et al. TFEB inhibits endothelial cell inflammation and reduces atherosclerosis. Sci Signal 10 (2017).

68. Justicia, C., Gabriel, C. & Planas, A.M. Activation of the JAK/STAT pathway following transient focal cerebral ischemia: signaling through Jak1 and Stat3 in astrocytes. Glia 30, 253–270 (2000).

69. You, T., et al. IL-17 induces reactive astrocytes and up-regulation of vascular endothelial growth factor (VEGF) through JAK/STAT signaling. Sci Rep 7, 41779 (2017).

70. Camporeale, A., et al. STAT3 activity is necessary and sufficient for the development of immune-mediated myocarditis in mice and promotes progression to dilated cardiomyopathy. EMBO Mol Med 5, 572–590 (2013).

71. Caston, R.A., et al. Combined inhibition of Ref-1 and STAT3 leads to synergistic tumour inhibition in multiple cancers using 3D and in vivo tumour co-culture models. J Cell Mol Med 25, 784–800 (2021).

72. Kun, L., et al. CRISPRi screens in human astrocytes elucidate regulators of distinct inflammatory reactive states. bioRxiv (2021).

73. Propson, N.E., Roy, E.R., Litvinchuk, A., Köhl, J. & Zheng, H. Endothelial C3a receptor mediates vascular inflammation and blood-brain barrier permeability during aging. J Clin Invest 131 (2021).

74. Zamanian, J.L., et al. Genomic analysis of reactive astrogliosis. J Neurosci 32, 6391–6410 (2012).

75. Fellmann, C., et al. An optimized microRNA backbone for effective single-copy RNAi. Cell Rep 5, 1704–1713 (2013).

76. Deverman, B.E., et al. Cre-dependent selection yields AAV variants for widespread gene transfer to the adult brain. Nat Biotechnol 34, 204–209 (2016).

77. Hasel, P., Rose, I.V.L., Sadick, J.S., Kim, R.D. & Liddelow, S.A. Neuroinflammatory astrocyte subtypes in the mouse brain. Nat Neurosci (2021).

78. Foo, L.C., et al. Development of a method for the purification and culture of rodent astrocytes. Neuron 71, 799–811 (2011).

79. Steiner, O., et al. Differential roles for endothelial ICAM-1, ICAM-2, and VCAM-1 in shear-resistant T cell arrest, polarization, and directed crawling on blood-brain barrier endothelium. J Immunol 185, 4846–4855 (2010).

80. Leow-Dyke, S., et al. Neuronal Toll-like receptor 4 signaling induces brain endothelial activation and neutrophil transmigration in vitro. J Neuroinflammation 9, 230 (2012).

81. Yenari, M.A., Xu, L., Tang, X.N., Qiao, Y. & Giffard, R.G. Microglia potentiate damage to blood-brain barrier constituents: improvement by minocycline in vivo and in vitro. Stroke 37, 1087–1093 (2006).

82. Toral-Rios, D., et al. Activation of STAT3 Regulates Reactive Astrogliosis and Neuronal Death Induced by AβO Neurotoxicity. Int J Mol Sci 21 (2020).

83. Ceyzériat, K., et al. Modulation of astrocyte reactivity improves functional deficits in mouse models of Alzheimer’s disease. Acta Neuropathol Commun 6, 104 (2018).

84. Rakers, C., et al. Stroke target identification guided by astrocyte transcriptome analysis. Glia 67, 619–633 (2019).

85. Priego, N., et al. STAT3 labels a subpopulation of reactive astrocytes required for brain metastasis. Nat Med 24, 1024–1035 (2018).

86. Toledo, J.B., et al. Contribution of cerebrovascular disease in autopsy confirmed neurodegenerative disease cases in the National Alzheimer’s Coordinating Centre. Brain 136, 2697–2706 (2013).

87. Serrano-Pozo, A., et al. Examination of the clinicopathologic continuum of Alzheimer disease in the autopsy cohort of the National Alzheimer Coordinating Center. J Neuropathol Exp Neurol 72, 1182–1192 (2013).

88. Mucke, L., et al. Astroglial expression of human alpha(1)-antichymotrypsin enhances alzheimer-like pathology in amyloid protein precursor transgenic mice. Am J Pathol 157, 2003–2010 (2000).

89. Abraham, C.R., Selkoe, D.J. & Potter, H. Immunochemical identification of the serine protease inhibitor alpha 1-antichymotrypsin in the brain amyloid deposits of Alzheimer’s disease. Cell 52, 487–501 (1988).

90. Ma, J., Yee, A., Brewer, H.B., Das, S. & Potter, H. Amyloid-associated proteins alpha 1-antichymotrypsin and apolipoprotein E promote assembly of Alzheimer beta-protein into filaments. Nature 372, 92–94 (1994).

91. Zhao, N., et al. Alzheimer’s Risk Factors Age, APOE Genotype, and Sex Drive Distinct Molecular Pathways. Neuron 106, 727–742.e726 (2020).

92. Bell, R.D., et al. Apolipoprotein E controls cerebrovascular integrity via cyclophilin A. Nature 485, 512–516 (2012).

93. Montagne, A., et al. APOE4 leads to blood-brain barrier dysfunction predicting cognitive decline. Nature 581, 71–76 (2020).

94. Clarke, L.E., et al. Normal aging induces A1-like astrocyte reactivity. Proc Natl Acad Sci U S A 115, E1896–E1905 (2018).

95. Fissolo, N., et al. CSF SERPINA3 Levels Are Elevated in Patients With Progressive MS. Neurol Neuroimmunol Neuroinflamm 8 (2021).

96. Williamson, M.R., Fuertes, C.J.A., Dunn, A.K., Drew, M.R. & Jones, T.A. Reactive astrocytes facilitate vascular repair and remodeling after stroke. Cell Rep 35, 109048 (2021).

97. Kumar, K.K., et al. Cellular manganese content is developmentally regulated in human dopaminergic neurons. Sci Rep 4, 6801 (2014).

98. Chambers, S.M., et al. Highly efficient neural conversion of human ES and iPS cells by dual inhibition of SMAD signaling. Nat Biotechnol 27, 275–280 (2009).

99. Chen, C., et al. Role of astroglia in Down’s syndrome revealed by patient-derived human-induced pluripotent stem cells. Nat Commun 5, 4430 (2014).

100. Ritchie, M.E., et al. limma powers differential expression analyses for RNA-sequencing and microarray studies. Nucleic Acids Res 43, e47 (2015).

101. Wang, J., Vasaikar, S., Shi, Z., Greer, M. & Zhang, B. WebGestalt 2017: a more comprehensive, powerful, flexible and interactive gene set enrichment analysis toolkit. Nucleic Acids Res 45, W130–W137 (2017).

102. von Mering, C., et al. STRING: known and predicted protein-protein associations, integrated and transferred across organisms. Nucleic Acids Res 33, D433–437 (2005).

103. Wheeler, M.A., et al. Environmental Control of Astrocyte Pathogenic Activities in CNS Inflammation. Cell 176, 581–596.e518 (2019).

