## Supplemental information for "Reactive astrocytes transduce blood-brain barrier dysfunction through a TNFα-STAT3 signaling axis and secretion of alpha 1-antichymotrypsin"

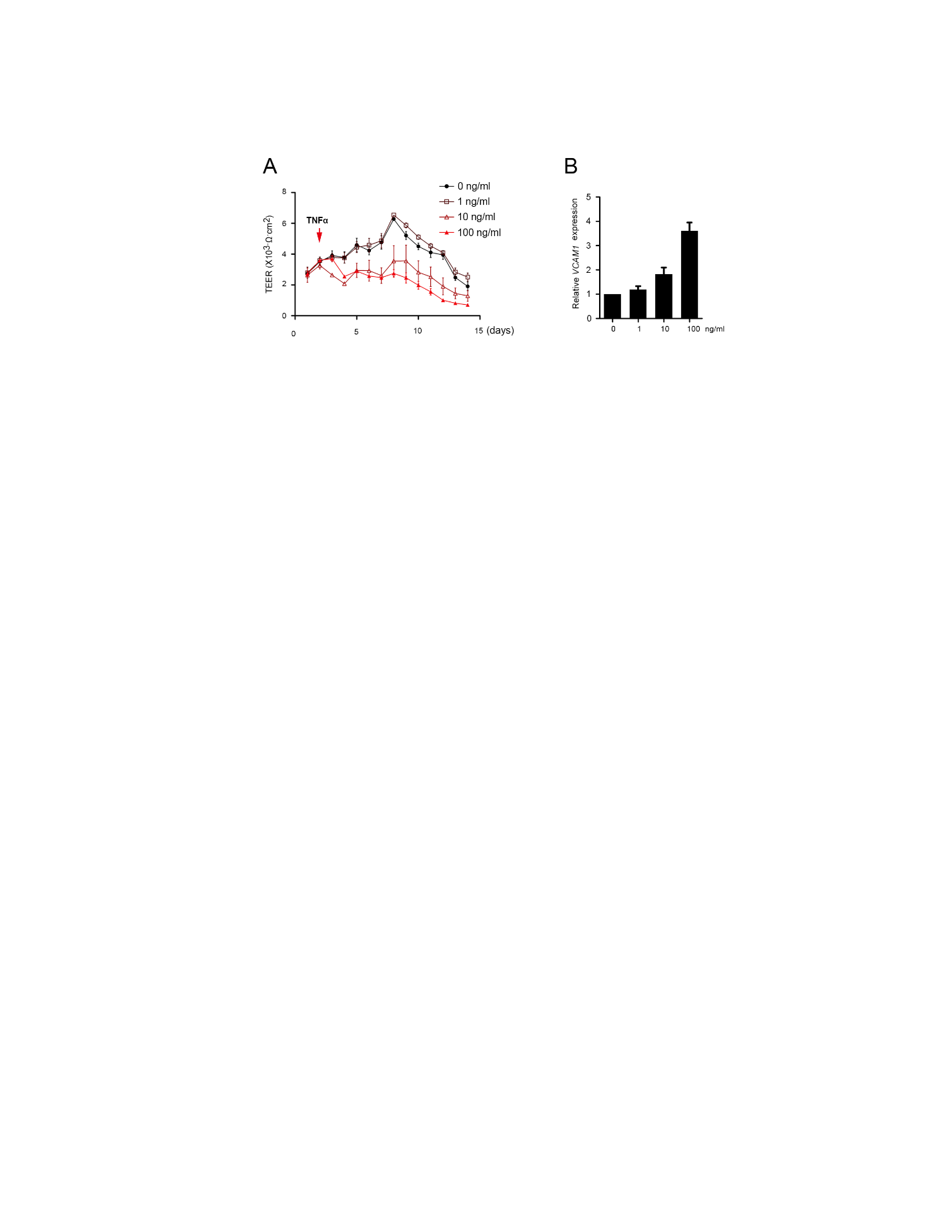

**Supplementary Figure 1. Dose-dependent effects of TNFα on cocultures of iPSC-derived BMEC-like cells and astrocytes.**

(A) Representative TEER values in BMEC-like cells as a function of TNFα dose. Data points represent mean ± SEM from duplicate Transwell filters per condition. Trends were confirmed across biological n=3.

(B) *VCAM1* expression in BMEC-like cells at day 14 as a function of TNFα dose. Data are normalized to the untreated control and graphed as mean ± SEM from n=3 biological replicates.

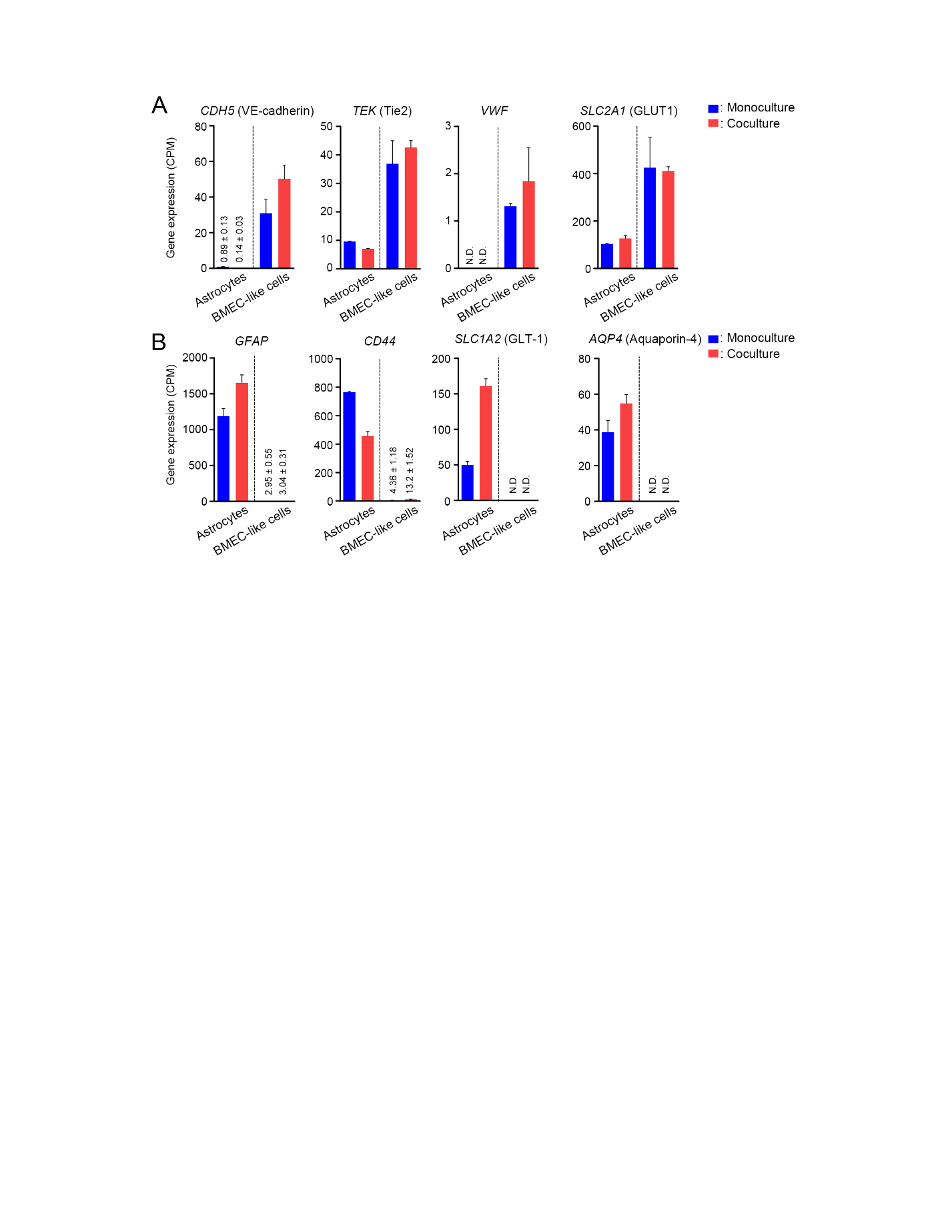

**Supplementary Figure 2. Validation of astrocytic and endothelial cell identity by cell-type-specific gene expression.**

Contrasted expression of endothelial-enriched (panel A) and astrocyte-enriched (panel B) genes in astrocyte and BMEC-like cells in monoculture or coculture. Data are graphed as mean ± SEM from n=3 biological replicates. N.D., not detected.

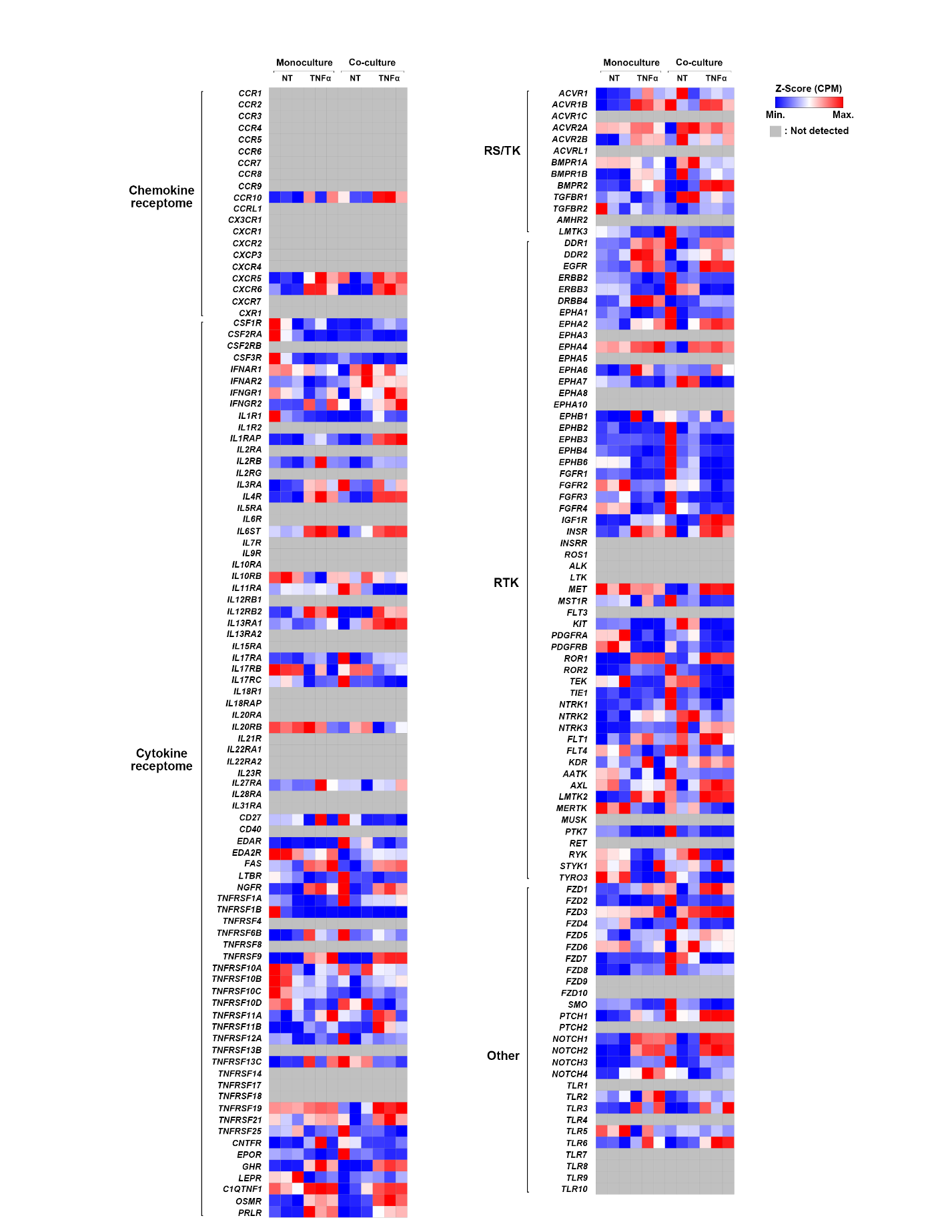

**Supplementary Figure 3. Receptome gene classifications in BMEC-like cells.**

Heat map showing gene expression of 194 human transmembrane receptors across all BMEC-like cell conditions. The gene sets are based on Kang et al., 2013^1^. The color of the heat map represents the CPM value for each gene.

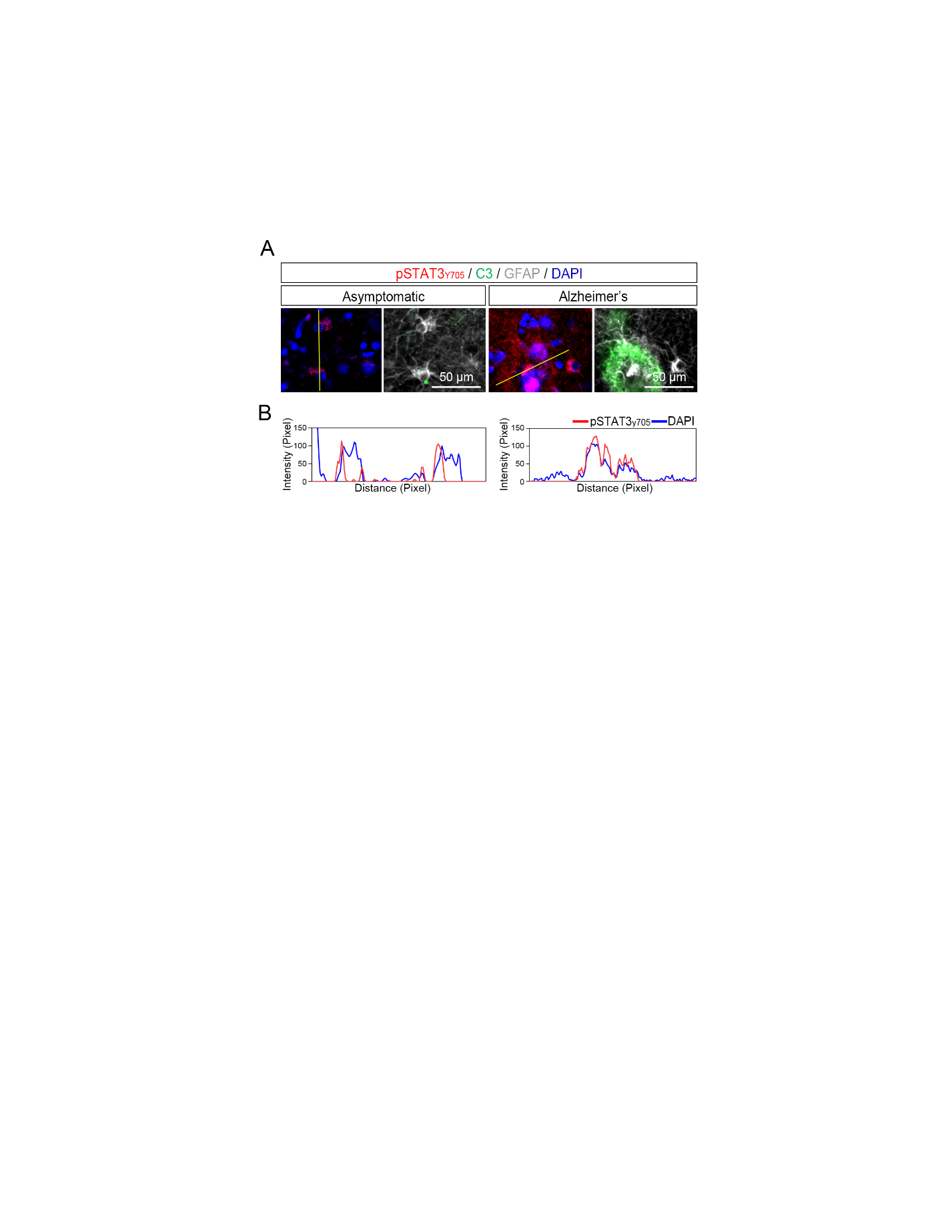

**Supplementary Figure 4. Intensity profile analysis for localization of pSTAT3_Y705_ in human brain tissue.**

(A) Representative images of pSTAT3_y705_, C3, and GFAP expression in an asymptomatic patient (left) versus a patient diagnosed with Alzheimer’s disease (right).

(B) Graphs showing one-dimensional intensity profile corresponding to each yellow line in panel A. Images were analyzed using ImageJ software.

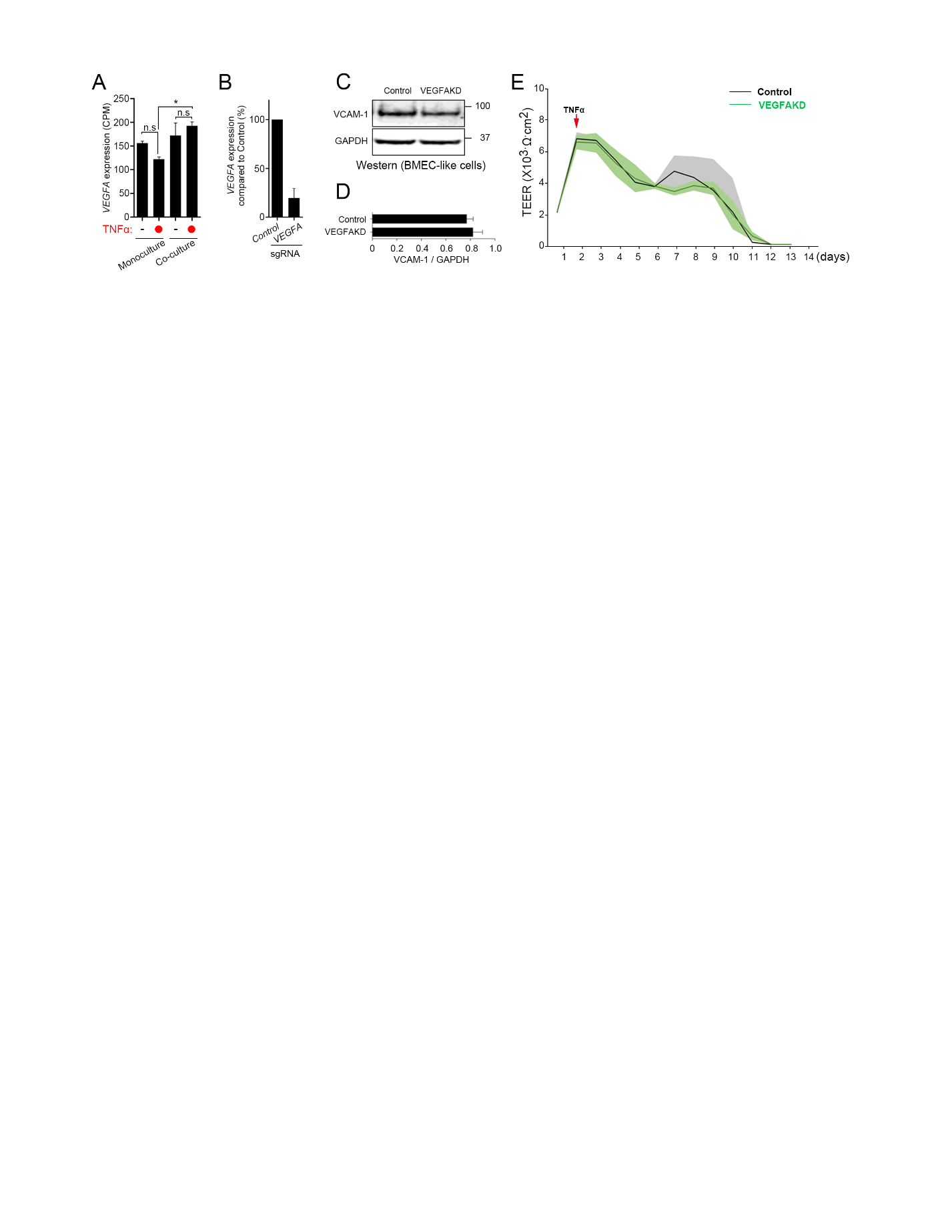

**Supplementary Figure 5. Effect of astrocytic *VEGFA* knockdown on BBB properties.**

(A) Expression levels (CPM) of *VEGFA* in across all astrocyte conditions. Data are graphed as mean ± SEM from n=3 biological replicates. One-way ANOVA with Tukey’s post hoc test: *, p<0.05.

(B) Relative *VEGFA* expression in CRISPRi-astrocytes transduced with sgRNA targeting *VEGFA*. Data are presented as mean ± SEM from n=3 biological replicates.

(C-D) Representative western blot (panel C) and quantification (panel D) of VCAM-1 expression in BMEC-like cells cocultured with CRISPRi-astrocytes*.* Samples were isolated on day 14 of coculture. Data are presented as mean ± SEM from n=3 biological replicates.

(E) TEER values in BMEC-like cells in coculture with CRISPRi-astrocytes transduced with sgRNAs. Data are presented as continuous means ± shaded SEMs aggregated from n=3 biological replicates per condition.

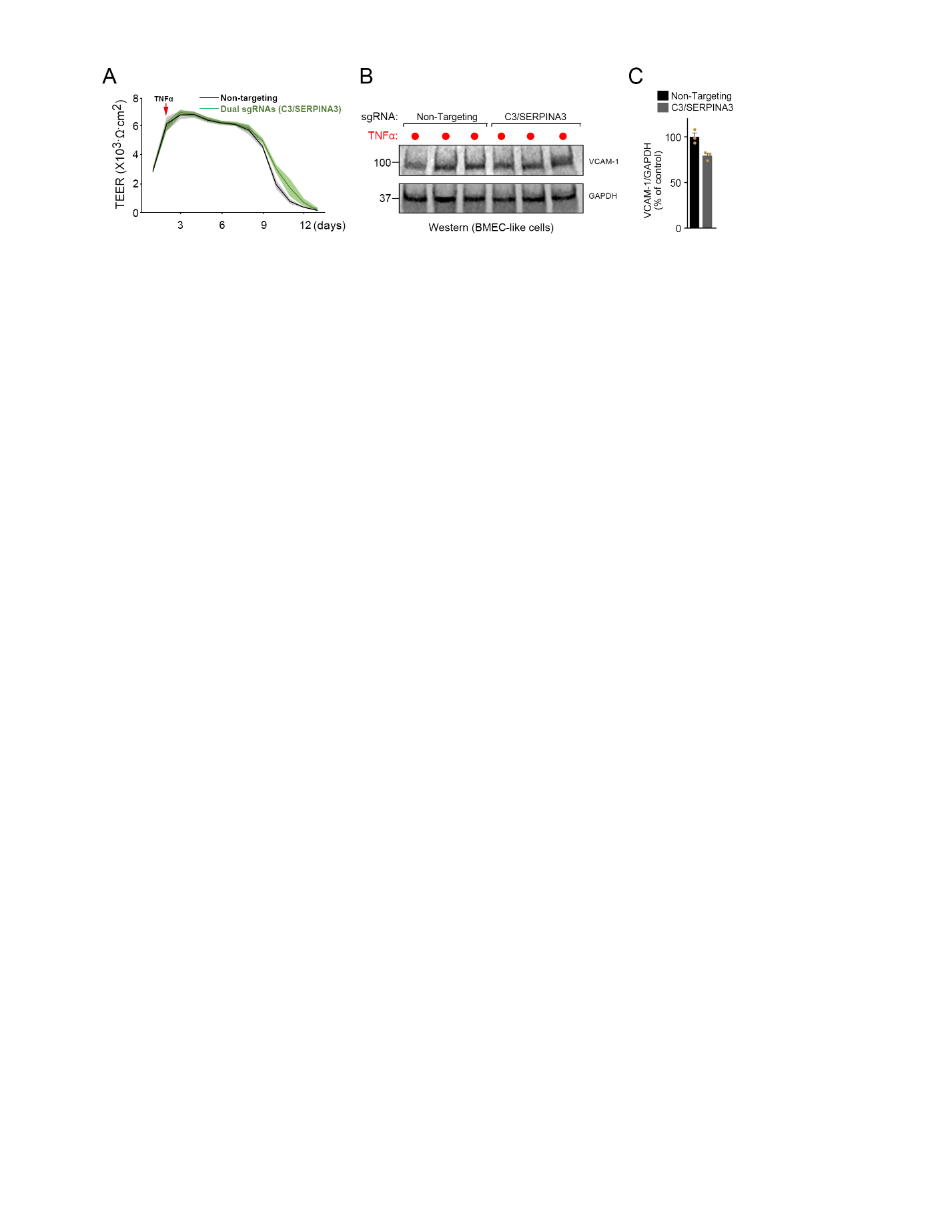

**Supplementary Figure 6. Effect of simultaneous *SERPINA3* and *C3* knockdown on BBB properties.**

(A) TEER values in BMEC-like cells in coculture with CRISPRi-astrocytes transduced with dual sgRNAs targeting *C3* and *SERPINA3*. Data are presented as continuous means ± shaded SEMs aggregated from n=3 biological replicates per condition.

(B-C) Representative western blot (panel B) and quantification (panel C) of VCAM-1 expression in BMEC-like cells cocultured with CRISPRi-astrocytes. CRISPRi-astrocytes were transduced with non-targeting sgRNA or dual sgRNAs targeting C3/SERPINA3. Samples were isolated on day 14 after coculture. GAPDH was used as a loading control. Quantification of western blot is graphed in panel C as mean ± SEM from n=3 biological replicates.

**Supplementary references**

1.	Kang, B.H., Jensen, K.J., Hatch, J.A. & Janes, K.A. Simultaneous profiling of 194 distinct receptor transcripts in human cells. *Sci Signal* **6**, rs13 (2013).
